# Genome evolution in an agricultural pest following adoption of transgenic crops

**DOI:** 10.1101/2020.10.05.326801

**Authors:** Megan L. Fritz, Kelly A. Hamby, Katherine Taylor, Alexandra M. DeYonke, Fred Gould

## Abstract

Replacement of synthetic insecticides with transgenic crops for pest management has been both economically and environmentally beneficial. These benefits have often eroded as pests evolved resistance to transgenic crops, but a broad understanding of the timing and complexity of adaptive changes which lead to field-evolved resistance in pest species is lacking. Wild populations of *Helicoverpa zea*, a major lepidopteran crop pest and the target of transgenic Cry toxin-expressing cotton and corn, have recently evolved widespread, damaging levels of resistance. Here, we quantified patterns of genomic change in wild *H. zea* collected between 2002 and 2017 when adoption rates of Cry-expressing crops expanded in North America. Using a combination of genomic approaches, we identified significant temporal changes in allele frequency throughout the genomes of field-collected *H. zea*. Many of these changes occurred concurrently with increasingly damaging levels of resistance to Cry toxins between 2012 and 2016, in a pattern consistent with polygenic selection. Surprisingly, none of the eleven previously described Cry resistance genes showed signatures of selection in wild *H. zea*. Furthermore, we observed evidence of a very strong selective sweep in one region of the *H. zea* genome, yet this strongest change was not additively associated with Cry resistance. This first, whole genome analysis of field-collected specimens to study evolution of Cry resistance demonstrates the potential and need for a more holistic approach to examining pest adaptation to changing agricultural practices.

**Significance Statement:** Evolution of pest resistance to management approaches in agricultural landscapes is common and results in economic losses. Early detection of pest resistance evolution prior to significant crop damage would benefit the agricultural community. It has been hypothesized that new genomic approaches could track molecular signals of emerging resistance problems and trigger efforts to pre-empt widespread damage. We tested this hypothesis by quantifying genomic changes in the pest *Helicoverpa zea* over a 15 year period concurrent with commercialization and subsequent loss of efficacy of transgenic Bt-expressing crops. Our results demonstrate the complex nature of evolution in agricultural ecosystems and provide insight into the potential for and pitfalls associated with use of genomic approaches for resistance monitoring. We discuss approaches for improvement.

## Introduction

The study of rapid evolutionary responses to natural selection has gained momentum in the past decade, and along with it recognition of the relevance of pest adaptation to control measures (Reznick et al. 2019). Melander published the first observation of insecticide resistance evolution in 1914 (Melander 1914), and since then, pests and weeds have adapted to myriad chemical, biological, and cultural control measures (Gould 1991, Palumbi 2001, Délye et al. 2013, Gould et al. 2018). While there has been an emphasis on finding single genetic loci that confer resistance to pesticides, recognition that surviving insecticide or herbicide applications could involve a suite of genomic changes has increased (Baucom 2019). Yet a clear understanding of the timing and complexity of changes associated with adaptation to agricultural ecosystems is lacking (Leon et al. 2020).

The bacterium, *Bacillus thuringiensis* (Bt), produces one or more crystalline (Cry) proteins that are each toxic to a narrow range of insect pests when ingested. Genes encoding Cry toxins were engineered into cultivated crops to reduce pest damage, producing so-called Bt crops, a major innovation in agricultural pest management that has been adopted around the globe (NASEM 2016). The economic and environmental benefits of this technology are significant: they can replace heavily used synthetic insecticides, which have damaging off target effects on pollinators and natural enemies, and have contributed to area wide pest suppression in non-Bt crops (Hutchison et al. 2010, NASEM 2016, Dively et al. 2018). A number of pest species have evolved resistance to one or more Cry toxins and there is concern about the durability of these transgenic crops (Tabashnik and Carrière 2017). Resistance management strategies have been implemented effectively for a few major pest species with high susceptibility to the toxins (NASEM 2016). Unfortunately, these measures have been inadequate for species where Cry toxins cause only moderate levels of mortality, providing opportunities for the evolution of resistance (Tabashnik and Carrière 2017).

Discovery of resistance-conferring genes has been of long-standing interest, in part due to expectations that this knowledge could lead to molecular diagnostic approaches that facilitate detection of early resistance evolution in wild pest populations (US-EPA 1998, US-EPA 2018). Molecular signals of resistance could then serve as triggers for regulatory actions that would minimize resistance risk and preserve environmentally sustainable technologies, like Cry toxins (US-EPA 1998). Genes discovered to have major impacts on Cry tolerance are being used to a limited extent for molecular resistance monitoring (Morin et al. 2004, Gahan et al. 2007, Zhang et al. 2013, Jin et al. 2018). To date, there are at least 11 such genes in the Lepidoptera (Wu 2014, Lin 2019), many of which were discovered in laboratory-selected populations and from screening wild populations for single genes with major effects on resistance (Morin et al. 2003, Gahan et al. 2001, Zhang et al. 2012, Jin et al. 2018). The question of whether molecular monitoring of these genes would effectively track resistance evolution in wild populations remains unresolved.

*Helicoverpa zea* is a migratory pest species (Gould et al. 2002) which comprises a single panmictic population in North America (Seymour et al. 2016). It has been controlled with Bt crops since 1996, when corn and cotton expressing Cry1A became commercially available. In 2007, corn and cotton cultivars producing both Cry1A and Cry2A were introduced. Since their commercialization, North American adoption rates of Cry-expressing corn and cotton grew from 8 and 15% of corn and cotton acreage in 1997 to *ca*. 80% by 2012. This rapid expansion of Cry-expressing crops placed significant selection pressure on their target pests, and widespread, damaging levels of resistance to Cry-expressing crops by *H. zea* were documented by 2016 (Dively et al. 2016, Reisig et al. 2018, Kaur et al. 2019, Yang et al. 2019). Existing Bt corn and cotton cultivars do not produce a high dose of Cry toxin relative to what is required to kill heterozygote *H. zea* larvae with resistance alleles (Luttrell and Jackson 2012). This likely contributed to their recent resistant status (Tabashnik and Carrière 2017). Under these conditions, resistance to Cry toxins was predicted to be polygenic (Gould 1998, Tabashnik et al. 2004) and to arise from standing genetic variation (Roush and McKenzie 1987).

Between 2002 and 2017, we saved *ca*. 1000 adult moths in ethanol each year, and we used samples from these collections to track genomic changes in *H. zea* over time. Instead of simply monitoring for changes in previously identified candidate genes, we took a gene agnostic approach and scanned entire genomes for signatures of selection. Using two sequencing approaches, double-digest Restriction-site Associated DNA sequencing (ddRAD-seq) and whole genome resequencing, we identified multiple genomic regions with significant changes in allele frequency during a 15 year time period. Shifts in allele frequency were used to calculate the strength of selection imposed on wild *H. zea* in agricultural ecosystems. We also examined whether any of the 11 previously identified candidate Cry resistance genes showed evidence of strong shifts in allele frequency over time. Surprisingly, none of these genes implicated in Cry-toxin resistance underwent significant temporal changes, nor did genes associated with detoxification of synthetic insecticides or plant defensive compounds, as was observed in another species where Bt crops replaced insecticidal control (Fritz et al. 2018). Instead, we found multiple novel candidate genes within regions of greater than expected allele frequency change, and quantified the effect size of the region showing the greatest temporal change on Cry resistance.

This first whole genome analysis of field-collected specimens to study impacts of Bt crops demonstrates the potential and need for a more holistic approach to examining pest adaptation to changing agricultural practices. While monitoring previously identified candidate genes may prove useful in some cases, our results indicate that such an approach is likely to miss critical evolutionary events. Analysis of patterns of genomic changes in pest populations following the introduction of a new control tactic or tool could prove useful, but requires follow-up experimental confirmation to demonstrate clear, causal relationships among genes and adaptive phenotypes.

## Results

### Strong temporal ddRAD-seq allele frequency shifts

*H. zea* males were collected from five pheromone baited traps (Hartstack 1979) in Bossier Parish, LA (SI Appendix, Table S1), and pooled by collection date in the months of May through October, 2002 through 2017. Randomly sampled moths from a subset of years were selected for genetic analysis; 2002, 2007, 2012, as well as in 2016, when strong phenotypic resistance to Cry1Ab and increasing resistance to Cry2Ab was being documented (Dively et al. 2016). Two hundred and sixty-five specimens were prepared into ddRAD-seq libraries and sequenced according to Fritz *et al*. (2016, 2018). Following filtering and alignment to the 341 Mb *H. zea* genome, we identified 14,398 single nucleotide polymorphisms (SNPs) from 259 individuals with an average read count of 404,003 (SI Appendix, Figure S1). Nucleotide diversity (π), and heterozygosity (F) did not change between years (SI Appendix, Table S2), and overall genomic divergence was low according to Weir and Cockerham’s *F*_ST_ (*F*_ST_ < 0.004; SI Appendix, Table S3). This low divergence was also supported by a sequential k-means analysis, which showed that a single genetic cluster or population (k = 1) best described our ddRAD-seq genotyped *H. zea* (SI Appendix, Figure S2). Interestingly, genetic divergence that occurred between the 2012-2016 collection periods was at least 3 times higher (*F*_ST_ = 0.0025) than for any other sequentially sampled periods. Pairwise *F*_ST_ values were 0.0009 and 0.0002 for the 2002-2007 and 2007-2012 collection periods, respectively (SI Appendix,Table S3).

We used a discriminant analysis of principal components (DAPC) to further examine temporal changes in the genomic structure of wild *H. zea*. Initially, we identified ddRAD SNP combinations that clustered individuals into groups (k = 2-4) by maximizing between group variation and minimizing within group variation. Using these SNP combinations, we estimated posterior membership probabilities for each individual from our dataset from each collection year. When k was equal to 2, samples within each of the collection years 2002, 2007, and 2012 were assigned to both genetic clusters. In contrast, samples collected in 2016 were strongly biased toward a single genetic cluster (Figure 1), and this trend persisted regardless of the k-value used (SI Appendix, Figure S3).). Interestingly, the genotypic shift between 2012 and 2016 corresponded to significant increases in the mean area consumed/ear of Cry1Ab + Cry2Ab expressing sweet corn (from near 0 to *ca*. 3cm^2^ between 2010-2016) and the proportion of two-toxin expressing ears containing late instars (from near 0 to > 75%; Dively et al. 2016). We sequentially removed the top 1%, 2.5% and 5% of ddRAD SNPs that had the greatest influence on genetic clustering, and re-evaluated population membership probability (assuming k=2). From this we determined that up to 5% (n = 720) of ddRAD SNPs were responsible for this observed genotypic shift (Figure S4).

**Figure 1.**
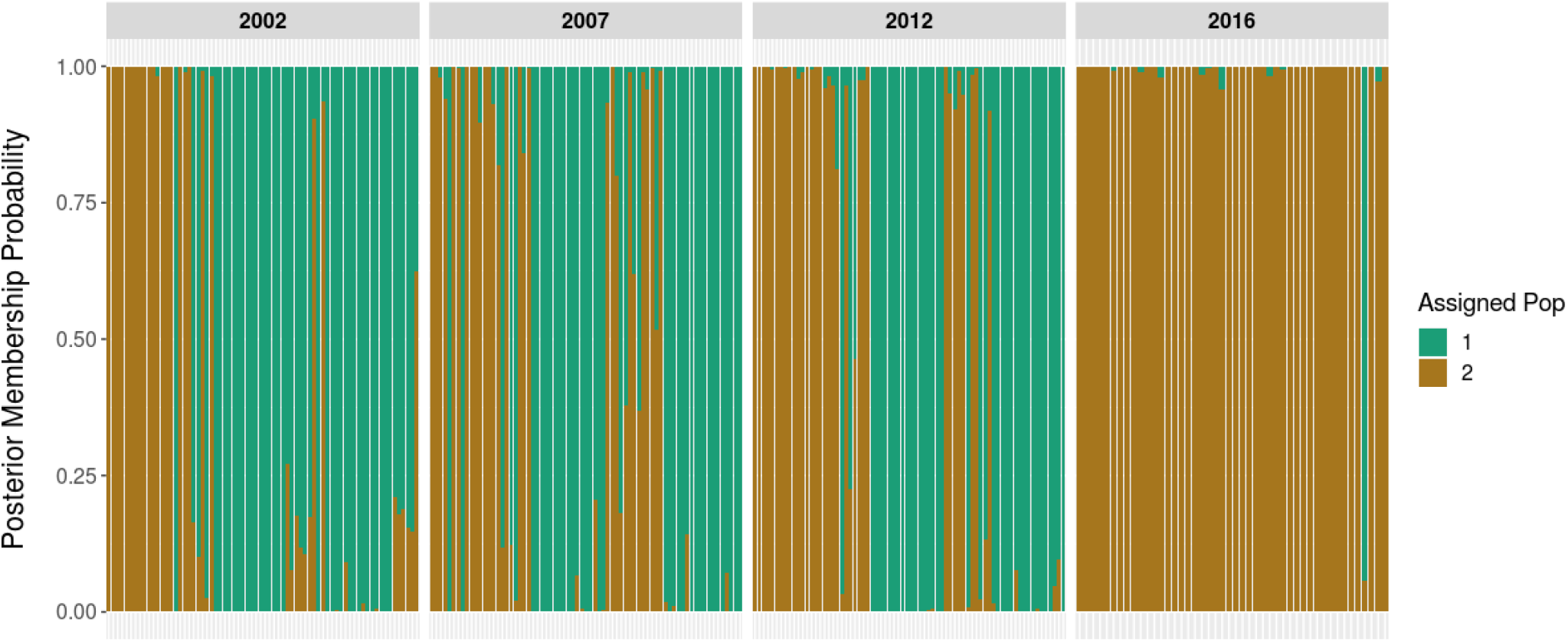
Posterior membership probabilities for *H. zea* individuals collected in Bossier Parish, LA, in 2002, 2007, 2012, and 2016. Genotypic clustering used ddRAD-seq generated SNP data and assumed a prior number of clusters (k) equal to two.

To identify which of the 14,398 ddRAD SNPs experienced greater than expected shifts in allele frequency over time, we conducted an *F*_ST_ outlier analysis (Whitlock and Lotterhos 2015). When we calculated genetic divergence (*F*_ST_) for each SNP across all 4 collection years, 53 had higher than expected divergence (Figure 2, File S1). Twenty-two of those 53 were among the top variance contributing SNPs that strongly influenced genetic clustering in our DAPC analysis. The 53 ddRAD SNP outliers were spread across 47 (of 2,975) unique scaffolds, and their among-year *F*_ST_ values ranged from 0.068 to 0.278.

**Figure 2.**
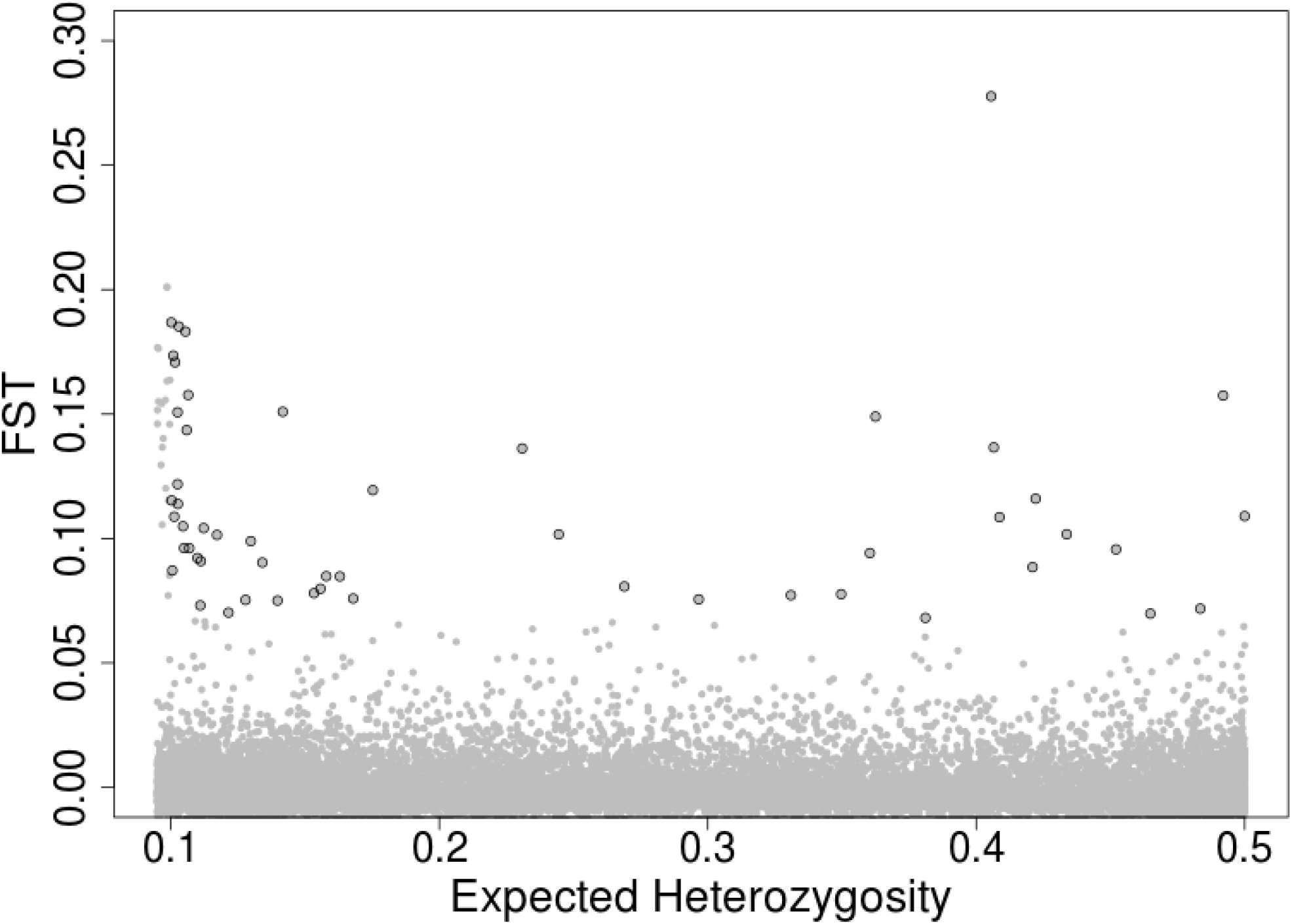
*F*_ST_ outlier analysis of *H. zea* collected from Bossier Parish, LA in the years 2002, 2007, 2012, and 2016. Each grey point represents one of the 14,398 SNPs examined in our total dataset. Those circled in black represent SNPs that were identified as outliers and indicate they were under directional selection.

### No significant allelic changes within candidate Bt resistance genes

We found ddRAD-seq markers within or near (< 56kb) 10 of the 11 previously identified candidate genes for Bt resistance, but none corresponded to outliers found over the time course of the collections (Table 1). To verify this result we conducted Illumina whole genome sequencing (WGS) of 35 *H. zea* males collected from the same location in LA in 2002 (n = 13), 2012 (n = 11), and 2017 (n = 11) (SI Appendix, Table S4). After genome alignment and filtering, this approach yielded 4,999,128 SNPs, and 2,622 fell within the 11 candidate Cry resistance genes. We used a 10kb sliding window-averaged Weir and Cockerham’s *F*_ST_ analysis and observed that *F*_ST_ values calculated for each gene ranged from −0.018 to 0.056 (Table 1; SI Appendix, Figure S5). The empirical threshold for statistically significant genetic divergence, generated according to Rubin et al. (2010), was 6 standard deviations (Z*F*_ST_ > 6) from the mean pairwise *F*_ST_ value in a window. None of the eleven candidate genes contained windows that reached our threshold for statistical significance of *F*_ST_ = 0.079 (Z*F*_ST_ > 6) per 10kb window. Taken together, these results indicated that the Cry resistance in *H. zea* was not caused by changes in the coding regions of any of these eleven candidate genes or their flanking cis-regulatory sequences.

**Table 1.** Candidate genes previously documented as being involved in Bt resistance in lab or field-collected Lepidopteran species, their best alignments to the *H. zea* genome. We calculated the average Weir and Cockerham’s *F*_ST_ value for SNPs within the gene-containing region of each scaffold. The number of SNPs used to calculate the average *F*_ST_ value can be found in parentheses. Whole genome sequencing samples from the years 2002 and 2017 were used for the comparison. We also report where the nearest SNP marker produced by ddRAD sequencing was found relative to each candidate gene and its *F*_ST_ value. When ddRAD SNPs were found within the gene, the average *F*_ST_ value across SNPs within the gene are reported. When the ddRAD SNP was found outside of the gene, the *F*_ST_ value for the nearest SNP is reported.

| Candidate Gene | Uniprot/ Genbank ID | Citation | *H. zea* scaffold | Gene Position (bp) | Avg $F_{ST}$ (Num SNPs) | Nearest ddRAD SNP (bp) | ddRAD Avg $F_{ST}$ (Num SNPs) |
| --- | --- | --- | --- | --- | --- | --- | --- |
| *alp* | ACP39715.1 | Jurat-Fuentes and Adang 2004 | KZ117832.1 | 1,946 - 16,347 | -0.011 (171) | NA | NA |
| *apn1* | Q11000.1 | Gill *et al.* 1995 | KZ118301.1 | 140,429 - 161,526 | -0.005 (353) | in gene | -0.003 (3) |
| *apn4* | AF378666.1 | Banks *et al.* 2001 | KZ118301.1 | 168,017 - 174,495 | 0.056 (170) | 22,238 | -0.005 (1) |
| *abcA2* | KP259911.1 | Tay *et al.* 2015 | KZ118207.1 | 171,103 - 192,554 | 0.002 (217) | in gene | 0.006 (2) |
| *abcC2* | GQ332571.1 | Gahan *et al.*<br>2010 | KZ118297.1 | 98,882 -<br>114,139 | -0.004<br>(100) | 3,178 | -0.004<br>(1) |
| *abcG1*<br>(white) | KM667971.1 | Guo *et al.*<br>2015a | KZ118424.1 | 2,680 -<br>24,508 | 0.015<br>(58) | 12,964 | 0.001 (1) |
| *calp* | AAK85198.1 | Gahan *et al.*<br>2001 | KZ117563.1 | 271,234 -<br>288,320 | 0.015<br>(201) | 56,926 | -0.007<br>(1) |
| *cad2* | AGG36450.2 | Zhang *et al.*<br>2017 | KZ118195.1 | 87,164 -<br>95,177 | -0.003<br>(44) | 20,535 | -0.003<br>(1) |
| *cad-86C* | *H. zea*<br>assembly v.<br>1.0<br>ID 537580 | Fritz *et al.*<br>2020 | KZ117463.1 | 550,935-603,715 | -0.008<br>(1079) | in gene | -0.004<br>(2) |
| *map4K4* | AKC96184.1 | Guo *et al.*<br>2015b | KZ118817.1 | 92,551 -<br>104,845 | -0.018<br>(65) | 35,125 | 0.005 (1) |
| *tspan1* | AYM54687.1 | Jin *et al.* 2018 | KZ118424.1 | 106,132 -<br>117,025 | -0.008<br>(164) | 27,211 | 0.003 (1) |

### Selection led to significant temporal divergence in novel genomic regions

Using our full WGS SNP dataset, we conducted pairwise analyses of genomic divergence between populations collected from 2002, 2012, and 2017 using 5, 10, 20 and 40 kb sliding window-averaged Weir and Cockerham’s *F*_ST_ along the length of the genome (SI Appendix, Figure S6). We report on the 40 kb sliding window analysis because it captured the major genomic changes that occurred for all sliding window sizes, but reduced the numbers of windows that appeared highly divergent due to few SNPs in the window. Thirty-six genomic windows on 11 different scaffolds had *F*_ST_ values that exceeded our empirical threshold for statistical significance of *F*_ST_ = 0.047 per 40 kb when 2002 and 2017 populations were compared (Figure 3). For paired comparisons of populations collected in 2002 and 2012, and 2012 and 2017 there were 55 and 30 genomic windows on 16 and 10 scaffolds with statistically significant genomic divergence, respectively (Figure 3; File S2).

**Figure 3.**
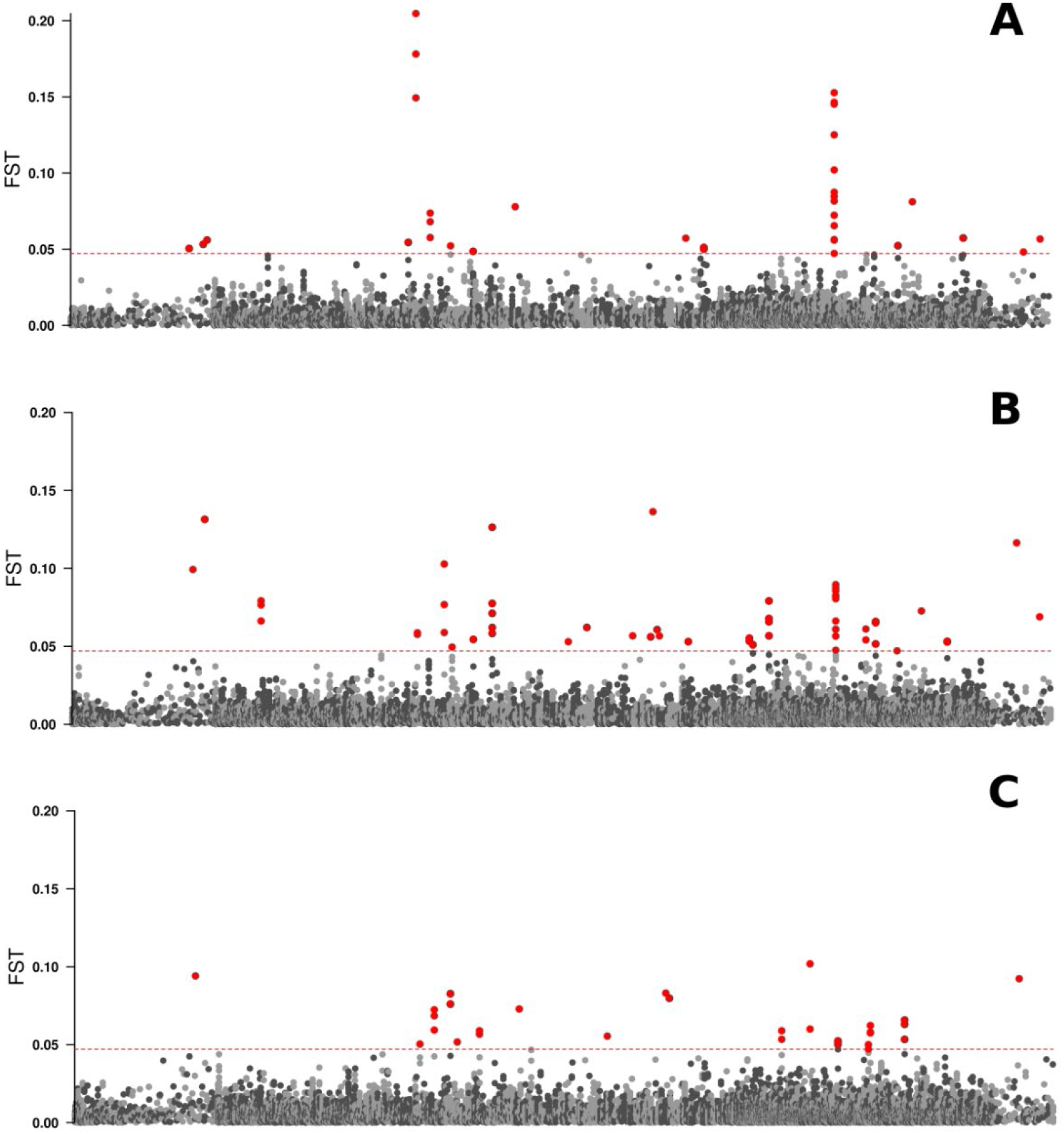
Manhattan plots of genomic divergence for individuals collected from pheromone-baited traps in Bossier Parish, LA in A) 2002 and 2017, B) 2002 and 2012, and C) 2012 and 2017. Alternating light and dark grey points demarcate averaged *F*_ST_ values for 40kb windows with a 10kb step for each of the 2,975 genomic scaffolds. Red points represent statistically significant genetic divergence.

Both genetic drift and selection influence the landscape of genomic divergence between populations. Therefore, we quantified the frequency with which our empirical threshold of *F*_ST_ = 0.047, calculated from the pairwise comparison of samples from 2002 and 2017, could be reached in a 40kb genomic window due to drift alone (Hudson 2002). Genotypic simulations were conducted for a single population sampled at two timepoints, but in the absence of selection. Allele frequency changes within 40kb windows were examined for multiple population sizes (N = 21,000 - 140,000). The proportion of genotypic simulations (of 20,000) that produced *F*_ST_ > 0.047 was the probability that our empirical threshold of *F*_ST_ = 0.047 could be exceeded due to genetic drift. Results from our simulations demonstrated this probability was very low (p = 0.0189 - 0) and negatively correlated with present day population size (SI Appendix, Figure S7 & Table S5).

Because the population sizes used in the simulations are likely underestimates of the effective population size of this migratory pest (Jones et al. 2019), it is highly improbable that our empirical *F*_ST_ threshold could have been exceeded by the action of drift. As a demonstration of its large effective population size, we collected fifth and sixth instar *H. zea* from ears of Obsession I (n = 8) and Cry-expressing Obsession II (n = 8) sweet corn grown in Central Maryland Research and Education Center (CMREC) demonstration plots. The genomes of these individuals, collected in September of 2017, were sequenced with Illumina short reads, and their genotypes compared to samples collected in Bossier Parish, LA, in that same year. After genome alignment and filtering for quality and linkage disequilibrium, we examined 14,697 SNPs in the dataset, which yielded a genome-wide *F*_ST_ value of 0.0004. The bootstrapped 95% confidence intervals were (−0.0002, 0.0008), and their overlap with zero indicated that allele frequencies in the LA and MD populations did not significantly differ from one another.

### Identification of novel genes under selection

We examined the genes near regions of greatest temporal genomic divergence to determine whether any were associated with detoxification of plant-produced or human-applied toxicants. Our list of candidates included those from families of genes associated with Cry resistance, but also resistance to synthetic insecticides and xenobiotics. Eleven genes, including 3 peptidases, 4 choline/carboxylesterases, 2 cytochrome p450s, 1 ABC transporter, and 1 tetraspanin, were near or within regions of significant temporal genetic divergence. No annotated proteases, proteinases, cadherins, voltage-gated channels, alkaline phosphatases, or glutathione-S-transferases were found within 50 kb of regions with elevated divergence. Evidence from a recent study suggests that increasing adoption of Bt crops led to a reduction in the frequency of synthetic insecticide resistance-conferring alleles, likely due to a reduction in synthetic insecticide applications (Fritz et al. 2018). Therefore, we specifically examined two genes known to be involved in pyrethroid resistance. Neither the voltage-gated sodium channel gene (*vgsc*), nor a cytochrome p450 (*cyp337b3*) which underlie pyrethroid resistance in the close *H. zea* relatives, *Chloridea virescens* (previously *Heliothis virescens*; Taylor et al. 1993) and *Helicoverpa armigera* (Joußen et al. 2012), respectively, were near regions of elevated genomic divergence. All genes found within 50kb of genomic windows with higher than expected divergence can be found in File S3.

We reasoned that increasing adoption of Bt crops over time would result in strong, sustained selection, leading to significant allele frequency changes across multiple collection years. Therefore, we identified scaffolds with elevated temporal genomic divergence in more than one pairwise comparison. Five scaffolds showed signs of significant allele frequency change in at least two pairwise comparisons. These included scaffold 173, which contained a *venom dipeptidyl peptidase 4* - like gene (HzOG204148), and scaffold 569, which contained two *carboxypeptidase Q (cpq)* -like genes (HzOG204151 and HzOG204153). Scaffold 1964 contained an RNA-directed DNA polymerase (HzOG206500), and scaffold 65 had one genomic window of elevated divergence in multiple pairwise comparisons which was near 7 genes, including *tetraspanin (tspan) 68C* (HzOG211510) and a *krueppel*-like gene (HzOG211515). Finally, scaffold 1612 contained 2 genomic windows with elevated divergence across all three by-year comparisons, which overlapped with *cyp333b3* (HzOG200024). Of these five scaffolds, 569 and 1612, had levels of genetic divergence that greatly exceeded all others between 2002 and 2017 (*F*_ST_ > 0.089), and such divergence was never observed for genetic simulations in the absence of selection (SI Appendix, Table S5).

The genome assembly of *H. zea* is fragmented into 2,975 genomic scaffolds, which makes detecting the extent of a selective sweep challenging. Under conditions of strong selection, the footprint in the genome should be broad, and we reasoned there was potential for some of these temporally divergent scaffolds to be physically linked. We used tblastn to align the genes from the 5 scaffolds showing strong temporal divergence in multiple by-year comparisons to the *Bombyx mori* genome (ASM15162v1), and identified their relative positions assuming conserved macrosynteny among Lepidoptera (d’Alençon et al. 2010). Four of these five scaffolds were adjacent to one another and syntenic to *B. mori* chromosome 13, indicating that the breadth of the selective sweep extended for *ca*. 3 Mb (SI Appendix, Figure S8). Linkage disequilibrium within the putative sweep region decayed more slowly than both genome-wide average rates of decay, as well as decay along single, similarly-sized scaffolds (> 1Mb) without evidence of genomic divergence (SI Appendix, Figure S9). This region of the *B. mori* genome contained 134 unique genes, none of which were previously known to be involved in Cry resistance (File S4). Interestingly, we re-examined our ddRAD-seq dataset and detected two ddRAD-seq markers found within this 3 Mb genomic region (on scaffold 569), which were outliers according to OutFLANK. This further confirmed the significance of changes occurring in this genomic region, but with a much larger sample size (n > 45 per year).

The 3 Mb putative sweep region experienced the greatest temporal change in frequency according to our whole genome scan. Yet our DAPC analysis identified 720 ddRAD SNPs on 505 unique scaffolds which caused a major architectural shift in the genomes of *H. zea* collected between 2012 and 2016 (Figure 1; SI Appendix Figures S3 & S4). Of those 720 ddRAD SNPs, only 5 aligned to this 3 Mb region. This pattern, consistent with polygenic selection on standing variation, indicated that allele frequency changes at many loci are involved in *H. zea* adaptation over the study period. To test how and whether strong temporal genomic changes should be used for genomic monitoring, our further analyses focused on the 3 Mb putative sweep region due to the breadth and magnitude of the observed allele frequency changes.

### Temporal shifts in the frequency of non-synonymous mutations

Given the significant levels of genomic divergence near *cyp333b3* and the two *carboxypeptidase Q-like (*hereafter *cpq*) genes detected by ddRAD-seq and WGS datasets, as well as the putative functional roles of these genes in insect metabolism and detoxification, we further examined protein-coding changes at these genes over time. Based upon our temporal analysis, genomic divergence was strongest at the 3’ end of all three genes (SI Appendix, Figure S10). While we detected 50 and 40 SNPs within *cpq1* and *cpq2*, respectively, only 3 were associated with putative changes in amino acid sequence (Table 3). One transition from cytosine to thymine caused a putative alanine to valine substitution in *cpq1*, and over time, the cytosine declined significantly from 0.73 in 2002, to 0.32 and 0.14 in 2012 and 2017, respectively (Freeman-Halton Fisher’s exact p = 0.0001; Table 2). Two SNPs in *cpq2* showed modest changes in allele frequency over time, although not statistically significant when a Bonferroni-corrected *α* value was used to account for multiple comparisons (Table 2). One cytosine to thymine transition encoded an alanine to valine amino acid substitution at 152,595 bp, and a second occurred at 152,183 bp, encoding a threonine to isoleucine substitution. Finally, there were 28 SNPs present in the protein-coding sequence of *cyp333b3*, 10 of which produced putative amino acid changes. Five of these non-synonymous mutations underwent statistically significant changes over time (p < 0.003; Table 2).

**Table 2 -.** Putative amino acid substitutions (AA Sub) for two *carboxypeptidase Q* genes (*cpq1* and *cpq2*) and *cyp333b3*. We quantified frequency changes in the codon present in the reference genome (Ref Codon) for *H. zea* populations collected in 1998 (n chr = 40-52), 2002 (n chr = 26), 2012 (n chr = 22), and 2017 (n chr = 22). A Freeman-Halton extension of a Fisher’s exact test with a bonferroni correction (*α* = 0.006) determined statistical significance (*). *Cpq1* extended from 8583 to 10,070 bp on scaffold 569, and *cpq2* extended from 151,736 to 160,706 bp, while cyp333b3 extended from 32,819 to 39,355 bp on scaffold 1612. All three genes are transcribed from the reverse strand.

| Scaffold | Gene | SNP Pos<br>(bp) | AA Sub | Ref<br>Codon | 2002<br>freq | 2012<br>freq | 2017<br>freq | p-val |
| --- | --- | --- | --- | --- | --- | --- | --- | --- |
| KZ118395.1 | *cpq1* | 9,409 | A → V | GCT | 0.73 | 0.32 | 0.14 | 0.0001* |
| KZ118395.1 | *cpq2* | 152,183 | T → I | ATA | 0.83 | 0.73 | 1 | 0.022 |
| KZ118395.1 | *cpq2* | 152,595 | A → V | GTA | 0.84 | 0.77 | 1 | 0.049 |
| KZ117131.1 | *cyp333b3* | 35,685 | S → T | AGC | 0.65 | 0.32 | 0.18 | 0.0001* |
| KZ117131.1 | *cyp333b3* | 36,339 | K → N | TTT | 0.69 | 0.27 | 0.09 | 0.0003* |
| KZ117131.1 | *cyp333b3* | 36,579 | G → V | GGA | 0.69 | 0.27 | 0.09 | < 0.0001* |
| KZ117131.1 | *cyp333b3* | 36,959 | T → A | GCC | 0.62 | 0.95 | 1 | < 0.0001* |
| KZ117131.1 | *cyp333b3* | 37,490 | S → N | AGC | 0.73 | 0.32 | 0.09 | 0.003* |

We also compared the frequencies of these non-synonymous mutations in the LA and MD populations collected in 2017, and for all 8 SNPs, there was no statistically significant difference in allele frequency based on collection location (SI Appendix, Table S7). Therefore, if allele frequency changes in these genomic regions were adaptive, they are not locally isolated to LA, but instead, present throughout the Eastern United States. These results and the low overall *F*_ST_ value for the two-population comparison (reported above) support previous findings that *H. zea* is highly migratory with little population structure.

### *Strength of selection experienced by wild* H. zea

Using the temporal changes in allele frequency, we calculated the strength of selection required to produce these statistically significant protein-coding changes over time. The selection coefficient(*s*) gave the strength of selection against *q*, the declining allele, examined under 3 different dominance scenarios of *p*: complete dominance, no dominance, and recessiveness. For 5 of the 8 non-synonymous mutations (Table 2), *q* was the allele present in the reference genome, and in the remaining cases, *q* was the alternate allele. We calculated mean values of *s* from allele frequency changes which occurred during the 2002-2012 collection period, as well as the 2012-2017 collection period. Values of *s* varied according to selection period and dominance scenario (Figure 4), but most fell within the range of 0.03-0.17. When *p* was assumed to be recessive, stronger selection was required to produce the observed changes in allele frequency in the 2002-2012 collection period due to its low frequency (*ca*. 0.25 - 0.3). Yet when *p* was dominant, stronger selection was required to produce the observed changes in allele frequency between 2012-2017, due to its relatively high starting frequency (*ca*. 0.65-0.75). In the case of the SNP at 36,959 bp on scaffold 1612, it’s frequency (*p*) was already 0.95 in 2012. Assuming dominance of *p*, the strength of selection that would have been required to increase its frequency to 1 by 2017 was very high (*s* = 0.70), which increased the variance around the mean value of *s* (Figure 4).

**Figure 4 -.**
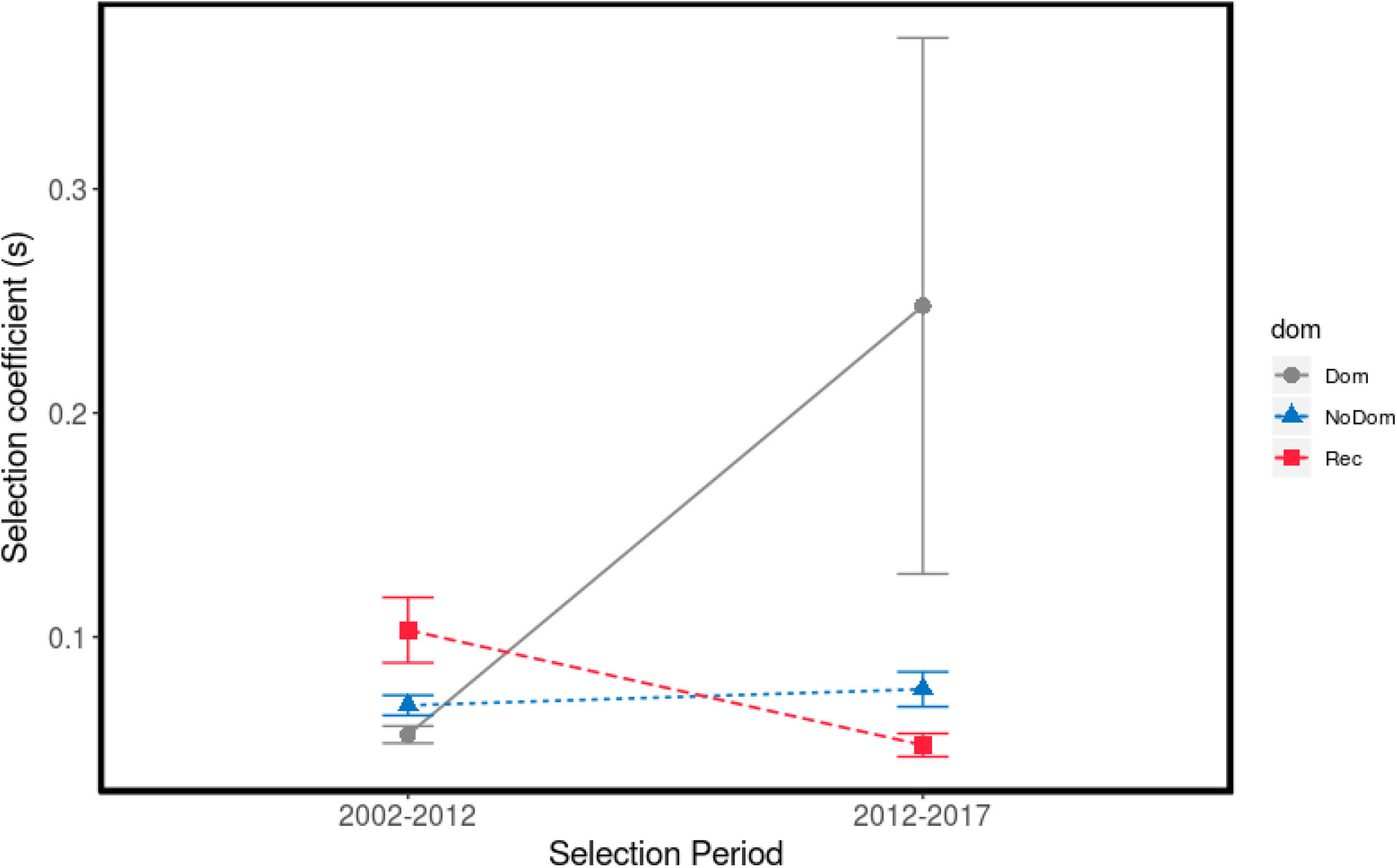
Values of *s*, the selection coefficient, required to produce the observed allele frequency changes at SNPs within protein-coding regions in field-collected *H. zea* during two selection periods: 2002-2012 and 2012-2017. Points represent the mean values (n = 6 SNPs), and error bars represent standard error of the mean.

### Frequency of non-synonymous substitutions prior to 2002

The intermediate frequency of the non-synonymous polymorphisms in 2002 compelled us to examine the frequencies of those polymorphisms in *H. zea* collected from our LA trapping sites in 1998, shortly after the release of Cry-expressing crops in 1996. Moths from 1998 had DNA quality that was too low for use in ddRAD-seq and WGS analyses, but short fragments could be PCR amplified. We therefore amplified and Sanger sequenced *ca*. 200bp DNA fragments spanning one non-synonymous polymorphism in each of *cpq1* and *cyp333b3* from 20-26 *H. zea* collected in 1998. The respective frequencies of the reference alleles for *cpq1* and *cyp333b3* were 0.93 and 0.90 in 1998 compared with 0.73 and 0.65 in 2002, and 0.14 and 0.18 in 2017 (Table 2) showing further evidence of genomic divergence at these sites. The change in allele frequency between 1998 and 2002 occurred prior to introduction of varieties with Cry2 genes.

### *Linkage of Cry resistance to* H. zea *Chr13*

To test whether this 3 Mb region with significant temporal genetic divergence was linked to Cry resistance, we bioassayed the progeny of two F2 genetic crosses between recently field-collected Cry resistant grandparents and Cry susceptible grandparents (Figure 5A & B). Cry resistant larval growth was 6 and 14 times greater than that of susceptibles when fed a diagnostic dose of diet containing Cry1Ab and Cry1A.105 + Cry2Ab2 sweetcorn leaf tissue, respectively, and these differences were statistically significant (Cry1Ab: df = 1, X^2^ = 314.57, p < 0.0001; Cry1A.105 + Cry2Ab2: df = 1, X^2^ = 409.97, p < 0.0001).

**Figure 5 –.**
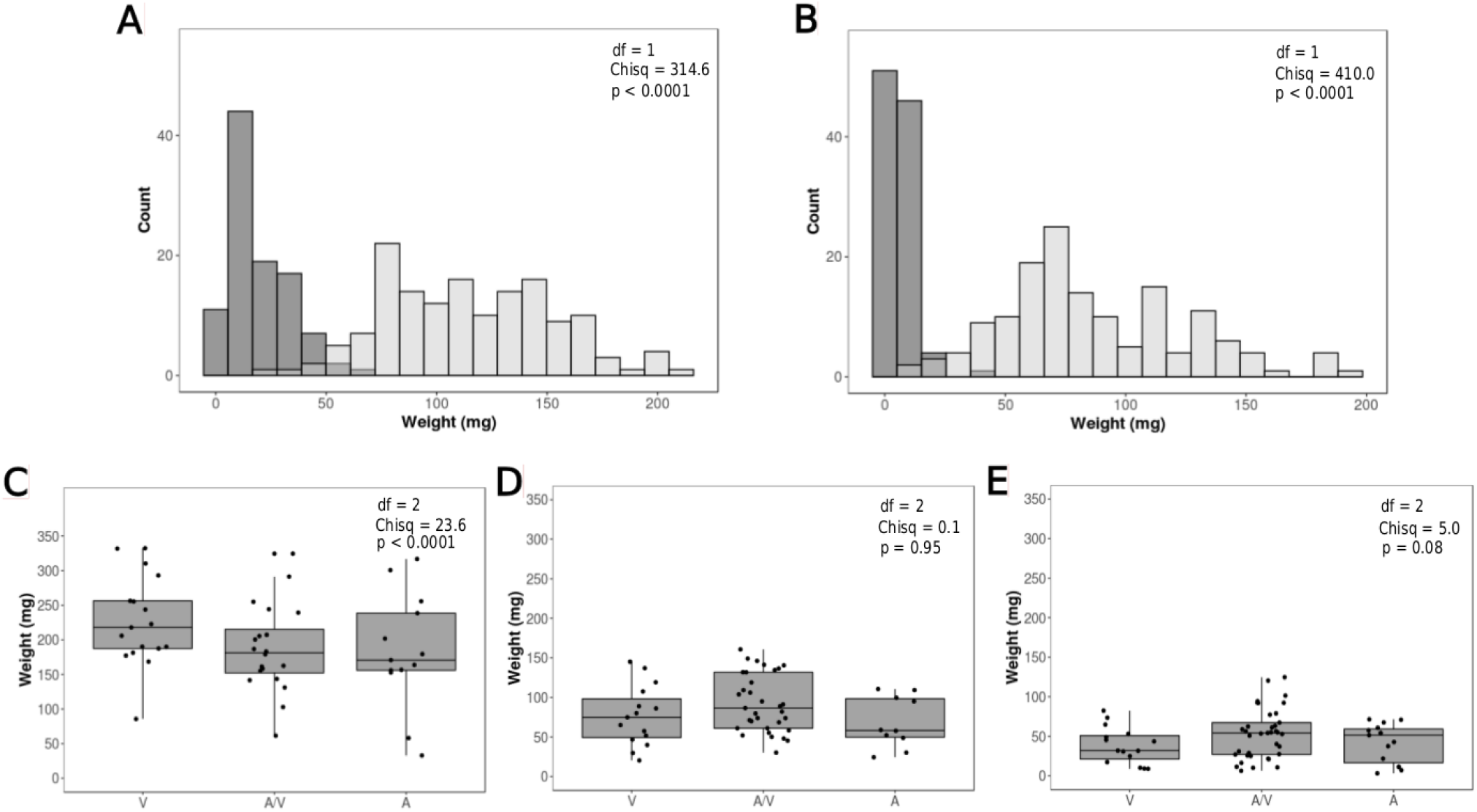
Distribution of parental populations and F2 larval growth phenotypes on Cry-treated diet. at a marker in the *cpq1* gene. Panels A and B show the distributions of larval weights for the susceptible (dark gray) and resistant (light gray) parental strains used for crosses after 7 days on diets containing Cry1Ab and Cry1A.105 + Cry2Ab2 expressing leaf tissue. Panels C, D, and E show the distributions of 7 day larval weights by *cpq1* marker genotype for progeny from two F2 mapping families. Grey boxes represent the upper and lower quartiles with the median (black horizontal line). Vertical lines show the minimum and maximum weights for each genotype, and points represent 7 day weights of each individual overlaid on the boxplots. V is the homozygous genotype putatively linked to Cry resistance, A/V represents heterozygotes, and A represents the homozygous genotype putatively linked to susceptibility.

Following 7 days of exposure to untreated, Cry1Ab, or Cry1A.105 + Cry2Ab2 treated diet, F2 larval growth phenotypes from each family were measured. Bioassayed individuals were genotyped by PCR and Sanger sequencing of fragments spanning two non-synonymous mutations in the *cyp333b3* and *cpq1* genes. Genotypes at these two loci were highly correlated within individuals due to physical linkage (Kendall’s tau = 0.989, z = 12.1, p < 0.0001), so further analysis focused only on *cpq1*. We postulated that individuals with the derived *cpq1* allele, which caused a Valine (V) substitution, should grow larger when placed on diet containing Cry-treated leaf tissue. In general, heterozygous F2s tended to grow larger on Cry1Ab and Cry1A.105 + Cry2Ab2 treated diet than did homozygotes (Figure 5D & E), and this trend was marginally significant for the F2s fed on two toxin diet (df = 2, χ^2^ = 5.03, p = 0.081). Surprisingly, F2 larvae fed upon a diet containing untreated leaf tissue grew significantly larger when they were homozygous for the derived allele (df = 2, χ^2^ = 23.64, p < 0.0001; Figure 5C). While the rapid allele frequency changes on chromosome 13 are fascinating, the environmental conditions that could have selected for both greater larval growth of homozygotes on untreated leaf tissue and marginally significant increases in growth of heterozygotes on treated diet, remain unresolved.

## Discussion

In this study, we compared the genomes of *H. zea* collected between 2002 and 2017 to quantify changes that occurred during a period of rapid adoption of a major agricultural innovation, Cry toxin-expressing crops, in the United States. We found substantial changes in the frequencies of SNPs throughout the *H. zea* genome that occurred in concert with the evolution of resistance to Cry toxins in field populations of *H. zea*. A major finding of our work is that none of the previously identified candidate Cry resistance genes in Lepidoptera showed signatures of selection in the field-collected *H. zea*, nor were genes in the strongest selective sweep region associated with resistance in an additive way. Instead, the timing and complexity of the genomic changes that occurred in *H. zea* were consistent with the predicted polygenic nature of Cry resistance in this species, and provide insight into how, when and whether genomic approaches could be used for resistance risk monitoring.

Over the past four decades there has been an emphasis on the search for single genes that confer resistance to control measures in insect pests and weeds, and we now have a catalogue of such genes (Ffrench-Constant 2013, Feyereisen et al. 2015, Délye et al. 2013, Heckel 2020). There are cases where a single gene diagnostic (Donnelly et al. 2016) or a genomic diagnostic targeted at multiple previously established candidate genes (Weetman et al. 2018) has been useful in monitoring for resistance, but recently there has been concern that these focused approaches could lead to complacency, as resistance evolves based on unexpected genes (Heckel 2020) or based on multiple genes (Délye et al. 2013, Van Leeuwen et al. 2020, Leon et al. 2020) that are already at low frequencies in pest or weed populations. Our work demonstrates that these concerns are appropriate. If any or all of the previously identified candidate genes had been used in a diagnostic for Cry resistance in *H. zea*, the diagnostic would have missed the coming problem.

While functional roles of specific genes that we identified have not yet been demonstrated, multiple *H. zea* genomic regions have changed dramatically since the introduction of Cry toxin-producing cotton and corn. The most compelling changes were found within a broad selective sweep on the *H. zea* homolog of *B. mori* chromosome 13. WGS and ddRAD-seq datasets both showed stronger than expected temporal genomic changes in a roughly 3 Mb region over the 15 year sampling period. When the results of the genome analyses were combined with PCR analysis of SNPs within candidate genes from 1998 samples from the same location, allele frequency changes of up to 0.79 occurred in these genes with metabolic functions or that are associated with xenobiotic resistance (Ward 1976, Guzov et al. 1998, Bown and Gatehouse 2004). Interestingly, two of these genes (*cpq1* and *cpq2*) were from families capable of binding Cry toxins in *H. armigera* (Da Silva et al. 2018).

Analysis of genetic crosses between resistant and susceptible strains found that this genomic region was not linked to Cry resistance in mendelian, or even additive way (Figure 5). Progeny on the diagnostic dose rarely reached the weights of their resistant parents (145-150mg), indicating that *H. zea* resistance is quantitative in nature. Yet those that did had at least one copy of the derived allele at *cpq1*. Furthermore, heterozygotes with one copy of the derived *cpq1* allele had marginally significant increases in larval growth on diet containing Cry1A.105 + Cry2Ab2 expressing leaf tissue, indicating a small, non-additive effect on resistance. Interestingly, individuals that were homozygous for the derived allele also had significantly greater larval growth on diet containing untreated leaf tissue, suggesting the benefits of genetic changes at this locus are not restricted to Cry resistance. This likely genotype by environment interaction may explain how the derived allele could reach high frequency in a way that selection for overdominance by Cry toxins alone could not. If the benefits of this allele were realized only in heterozygotes exposed to Cry toxins, selection should have maintained this allele at intermediate frequency. The functions of genes in the 3 Mb sweep region are not clear, particularly how or why they promote increased larval growth on non-expressing plant tissue. Yet the heterogeneous North American environment, with selective pressures from both Cry-expressing and non-expressing host plants of *H. zea*, provides a landscape with capacity to drive the derived allele to a frequency greater than 0.5. This finding highlights both the complex nature of Cry resistance in *H. zea*, as well as risks associated with genomic monitoring. Strong selective sweeps do not necessarily point to resistance genes with large effect sizes, and future studies should be directed at understanding what sweeps in regions with complex genotype by environment interactions can tell us about resistance risk.

We emphasize that we detected multiple regions of the genome that changed during the course of our sampling, a pattern consistent with polygenic adaptation from standing genetic variation. Our findings are consistent with a recent study of crosses of Cry2 resistant *H. zea* to a susceptible strain that found phenotypic variance fitting a polygenic model (Yang et al. 2020). In our study, strong polygenic change was most apparent between the years 2012 and 2016 (Figure 1) when widespread and damaging levels of resistance to Cry toxins were documented (Dively et al. 2016). Ongoing, experimental, quantitative genomic studies will shed light on which of these genomic regions contribute most substantially to Cry tolerance in field-collected larvae.

The use of genomic approaches to study rapid evolution in natural populations has gained momentum (*e.g*. Crossley et al. 2017, Fritz et al. 2018, Nosil et al. 2018, Weetman et al. 2018, Bi et al. 2019, Dayan et al. 2019, Laurentino et al. 2020). Yet their use for studying pest adaptation to agricultural systems has been limited to a few cases (reviewed in Pélissié et al. 2018). As we attempt to apply these approaches to a variety of agricultural systems, system- and method-specific issues will need to be addressed to ensure accuracy. Results from our experiments indicate that genomic approaches may hold promise for monitoring resistance risk in pest populations exposed to novel management practices, with several caveats for how we interpret our results.

One concern is that major genomic changes identified through genomic monitoring might not be due to the control tool being studied. Deployment of pest management innovations do not take place in the absence of other existing selection pressures in the environment. When transgenic cotton and corn were introduced into North American agricultural systems, the use of synthetic insecticides were reduced, though certainly not eliminated. In a previous study using ddRAD-seq, we found that another pest of cotton in Louisiana, *Chloridea virescens*, did not experience changes in the frequency of Bt resistance candidate genes. Rather, changes to the frequency of an allele that conferred resistance to pyrethroid insecticides were observed over a 15 year period (Fritz et al. 2018). Indeed, our data suggest that the greatest allele frequency changes in the *H. zea* genome were not solely associated with Cry resistance. Therefore, confirming the strength of association between any major genomic change with a resistance phenotype is critical.

This confirmation testing can be conducted using the types of simple crosses and PCR-based assays that we carried out in our study. Such assays can determine the importance of rapidly changing loci to resistance phenotypes and alleviate concerns regarding resistance risk. Conversely, knowledge of allele frequency changes and confirmation of their impact on field resistance could then be used to make predictions for future changes in resistance in field populations of pests. In cases similar to *H. zea* on Cry-expressing crops where the control tactic does not kill all pest individuals or has sublethal effects, collections from paired sentinel plots could also point to genomic regions associated with resistance. Such paired sentinel plots are already being used for determining the current extent of resistance, but typically monitor damage to the crop rather than the genetics of the pest (Dively et al. 2016, 2020). Focusing on damage and the economic impact of resistance does little to predict risk of resistance and then mitigate it. Once widespread damage has occurred, resistance alleles have likely already spread widely in the landscape. Instead, sampling and sequencing early instar larvae found in treated and untreated sentinel plots would allow researchers to quantify genomic change associated with the pest management tool compared to the population at large.

A second concern is that population demographic processes, including migration patterns and population expansion can result in the appearance of genome-wide shifts in allele frequency, which are not due to selection. When this occurs, the risk for genomic monitoring would be to falsely implicate a region of the genome in evolving resistance. We were fortunate to be studying a highly migratory species (Hartstack et al. 1982, Gould et al. 2002, Seymour et al. 2016) with little population structure over large geographic areas (Perera and Blanco 2011, Seymour et al. 2016). Our finding of negligible genomic differences between samples collected in 2017 from Louisiana and Maryland are consistent with results from these previous studies. Additionally, *H. zea* feeds on a number of host plants, and this seems to have resulted in the population densities remaining similar to those prior to the use of transgenic cotton and corn (Micinski et al. 2008). It is not surprising that with this species, measured overall genomic diversity remained constant. For a species that interbreeds over such a large area, population demographic processes are to result in detection of false positives. Had we been studying an insect or weed with strong local population structure, or one in which the control measure caused severe reduction and resurgence of the pest, our experimental approaches would have required additional careful consideration.

Finally, many so-called “mega-pest” lineages are thought to have genomes that already contain variants, which prime them for the evolution of resistance to novel pest control measures (Crossley et al. 2017, Pélissié et al. 2018). When selection is applied to these populations, they experience soft selective sweeps at genomic regions that are typically spread throughout the genome. Such signatures of polygenic selection in field-collected samples can be difficult to bioinformatically tease apart from the impacts of drift. Fortunately, new computational approaches which can better differentiate soft selective sweeps from background selection or other population demographic processes are becoming available (Schrider and Kern 2016), and some even detect the proportion of genome-wide changes in populations due to selection using temporal datasets (Buffalo and Coop 2019). Further development of bioinformatic approaches specifically aimed at detecting temporal genomic changes at both single loci and those spread throughout the genome will provide critical tools for resistance risk monitoring in agricultural systems.

The benefits of genomic monitoring of pest species go beyond identifying the mechanisms of resistance and applications for resistance risk mitigation. Adoption of genomic approaches for resistance monitoring will be crucial for addressing the decades-long debate on exactly how pest species rapidly adapt to management practices (Crow 1957). Even today, the extent to which a species response to selection is mono- or polygenic, from standing genetic variation or novel mutation remains unclear. Here we present evidence on the complex nature of genomic change during expansion of a major agricultural innovation, Cry-expressing crops, in one important pest species. Extension of genomic monitoring programs, accompanied by targeted experimental confirmation, to additional pest species on broad spatial and temporal scales could provide critical insight into the nature of adaptation by these species to human-imposed selection.

## Data Availability

Data for this study are available at: to be completed after manuscript is accepted for publication. All scripts and barcodes files used to perform these analyses can be found at https://github.com/mcadamme/FieldHz_Pop_Genomics.

## Acknowledgements

We thank David O’Brochta, Brad Coates, David Heckel, Bruce Tabashnik, and our anonymous reviewers for suggestions that improved our manuscript. This work was funded by USDA NIFA Biotechnology Risk Assessment Grants 2016-33522-25640 and 2019-33522-29992.

## Methods

### Insect collections for ddRAD-seq and WGS analyses

*H. zea* adults were collected by pheromone-baited trap bi-weekly from May through September in Bossier Parish, Louisiana (SI Appendix, Table S1) in 2002, 2007, 2012, 2016, and 2017. Late instar (5-6^th^) *H. zea* were also collected in 2017 at the University of Maryland CMREC farms in Prince George’s county, MD, and reared to adulthood prior to genomic analysis.

### DdRAD-seq library preparation

Specimens from the years 2002, 2007, 2012, and 2016 were prepared into double-digest RAD-seq (ddRAD-seq) libraries for population genomic analysis according to Fritz *et al*. (2016, 2018), and sequenced on 8 runs of an Illumina MiSeq.

### DdRAD-seq Bioinformatic Analysis

Merged, filter-trimmed reads were aligned to the *H. zea* genome (v. 1.0, NCBI Bioproject PRJNA378438; Pearce et al., 2017) with Bowtie2 (Langmead and Salzberg 2012). Genotypes were called with BCFtools (Danecek et al. 2016) and filtered to include SNPs that i) had a depth of coverage > 3, ii) were present in at least 75% of the 259 individuals in our filtered dataset, iii) had a minor allele frequency of 0.05, iv) had a maximum of 2 alleles, and v) were thinned to 1 per 200 bp to reduce the degree of linkage disequilibrium between SNPs on the same ddRAD-seq marker.

We calculated average nucleotide diversity (π) and heterozygosity (the inbreeding coefficient F) values for populations collected in each year, and examined genome-wide divergence for each pair of years using pairwise *F*_ST_ (Weir and Cockerham 1984). A k-means analysis in R (R core team 2008) modeled the number of putative populations or clusters (k = 1:10) into which the LA *H. zea* fit according to genotype (adegenet v. 2.1.1, Jombart et al. 2010, Jombart and Ahmed 2011). Using the three best-fitting numbers of clusters (k = 2:4) as priors, we applied a Discriminant Analysis of Principal Components (DAPC) in adegenet to identify groups of genetically similar *H. zea* individuals. Weir and Cockerham’s *F*_ST_ was calculated for each SNP to quantify genetic divergence across years and identify those with higher than expected divergence over time (OutFLANK v. 0.2, Whitlock and Lotterhos 2015). Locations of genes already implicated in Bt resistance in the *H. zea* genome were identified by blast, and we used custom scripts to identify the nearest ddRAD-seq marker to each (Table 1).

### WGS library preparation

Genomic DNAs from additional adults collected in 2002 (n= 13), 2012 (n = 11), and 2017 (n = 11) were prepared into two Illumina TruSeq LT libraries (Illumina, Inc. San Diego, CA), and sequenced on two Illumina NextSeq500 runs.

### WGS bioinformatic analysis

Following filter-trimming and reference genome alignment, called SNP genotypes were filtered to removed SNPs i) with a coverage depth < 3, ii) absent from 8 or more individuals sequenced, iii) with minor allele frequency of < 0.05, and iv) with > 2 alleles per locus. Temporal change at Bt resistance candidate genes was measured by pairwise *F*_ST_ averaged across SNPs found within each gene. We applied a sliding window analysis, where windows with an average *F*_ST_ value greater than 6 standard deviations from the mean (Z*F*_ST_ > 6) were considered to have undergone statistically significant divergence (Rubin et al. 2010).

Sliding window averaged *F*_ST_ values were also used to identify novel signatures of strong selection across the genome using 40kb windows with a 10kb step. Additional sliding window and step sizes were also examined (SI Appendix, Figure S2). Genotype simulations under neutral expectations using realistic population demographic conditions generated a distribution of *F*_ST_ values, which were compared against empirical *F*_ST_ values from our genome scan as a secondary approach to examining statistical significance (Hudson 2002; SI Appendix, Supplemental Methods). Results of our whole genome scan were compared to results from our ddRAD-enabled outlier analysis to determine which regions were detected in both datasets. We queried lists of genes within 50kb of each novel genomic window that underwent statistically significant temporal divergence to identify those in families associated with metabolism and detoxification (SI Appendix, Supplemental Methods). For the 5 most divergent scaffolds, we determined scaffold order and proximity by tblastn alignment of protein-coding regions to the *B. mori* genome, assuming conservation of macrosynteny among Lepidopteran genomes (d’Alençon et al. 2010).

### *Genome similarity between* H. zea *collected from MD and LA in 2017*

A global *F*_ST_ analysis compared *H. zea* collected at CMREC in Prince George’s county, MD, in 2017 to those collected in Bossier Parish, LA, in 2017, and overlap of the bootstrapped 95% confidence intervals with zero determined statistical significance.

### Gene-specific analyses

Temporal and spatial differences in the frequencies of non-synonymous substitutions within *cpq1, cpq2* and *cyp333b3* were analyzed with a Bonferroni-corrected two-sided Fisher’s exact test.

### Strength of selection

We calculated strength of selection (*s*) imposed upon *cpq1, cpq2*, and *cyp333b3* non-synonymous substitutions under 3 different dominance scenarios: dominance of the allele that was increasing in frequency, *p*, no dominance of *p*, and recessiveness of *p* (Falconer and Mackay 1996). Our model assumed 5 generations per year (Reay-Jones 2019) for each selection period: 2002-2012 and 2012-2017.

### Frequency of non-synonymous substitutions prior to 2002

Genomic DNA was isolated from *H. zea* collected in 1998 from Hartstack traps in Bossier Parish, LA, PCR-amplified using primers flanking non-synonymous mutations in *cpq1* and *cyp333b3* (SI Appendix, Table S8), and Sanger sequenced, allowing us to determine their frequency prior to 2002.

### Test for linkage

Using F2 progeny from crosses between Cry-tolerant field-collected and Cry susceptible laboratory-reared *H. zea* (Benzon Research Inc., Carlisle, PA), we measured 7-day larval growth phenotypes following exposure to Cry-treated and untreated diet (Dively et al. 2016; SI Appendix, Supplemental Methods). All F2s were genotyped at non-synonymous mutations in the *cyp333b3* and *cpq1* genes by PCR and Sanger sequencing, but due to linkage disequilibrium between markers, only *cpq1* was further analyzed. We used linear models to quantify the effect of *cpq1* genotype on 7 day larval weight following exposure to treated or untreated diets. Statistical significance was determined using model comparison by likelihood ratio test (SI Appendix, Supplemental methods).

