## Supplementary material for "Genome evolution in an agricultural pest following adoption of transgenic crops": SI_Appendix

**Supplemental Files**

**File S1** - DdRAD-seq SNPs showing significant genomic divergence among *H. zea* collected in 2002, 2007, 2012, and 2016.

**File S2** - Scaffolds containing 40kb genomic windows with higher than expected sliding window averaged $F_{ST}$ values for 3 by-year comparisons.

**File S3** - Genes within 50kb of scaffolds showing significant genomic changes over time. These changes were detected in *H. zea* adults collected in Bossier Parish, LA, in 2002, 2012, and 2017.

**File S4** - *Bombyx mori* genes found within the 3 Mb genomic region on chromosome 13 that is associated with the selective sweep in *H. zea*.

**Supplemental Tables**

**Table S1 –** GPS coordinates for pheromone-baited trapping locations in Bossier Parish, LA.

| **State** | **Trap No.** | **Latitude (deg N)** | **Longitude (deg W)** |
|---|---|---|---|
| LA | Hv_20 | 32° 25' 7.66" | 93° 38' 5.03" |
| LA | Hv_21 | 32° 24' 59.55" | 93° 38' 18.62" |
| LA | Hv_24 | 32° 24' 51.28" | 93° 38' 7.05" |
| LA | Hv_26 | 32° 24' 50.77" | 93° 38 '30.84" |
| LA | Hv_28 | 32° 25' 19.03" | 93° 38 '30.85" |

**Table S2** - Estimates of average nucleotide diversity ($\pi$)**,** heterozygosity (F), and their associated 2.5% and 97.5% quantiles across the 14,398 SNPs from our filtered ddRAD-seq dataset (n = 259 individuals).

| | **N** | $\pi$ | **2.5, 97.5% quantiles** | **F** | **2.5, 97.5% quantiles** |
|---|---|---|---|---|---|
| **2002** | 70 | 0.280 | 0.276, 0.283 | 0.190 | 0.152, 0.224 |
| **2007** | 71 | 0.280 | 0.277, 0.282 | 0.155 | 0.124, 0.175 |
| **2012** | 72 | 0.281 | 0.279, 0.284 | 0.167 | 0.146, 0.181 |
| **2016** | 46 | 0.286 | 0.284, 0.289 | 0.114 | 0.060, 0.160 |

**Table S3** - Pairwise genetic divergence between years at 14,398 SNPs generated by BCFtools across the *H. zea* genome according to Weir and Cockerham's $F_{ST}$. By-year comparisons with asterisks indicate statistically significant genetic divergence at the $p < 0.05$ level.

| | **2002** | **2007** | **2012** | **2016** |
|---|---|---|---|---|
| **2002** | - | - | - | - |
| **2007** | 0.0009* | - | - | - |
| **2012** | 0.0011* | 0.0002 | - | - |
| **2016** | 0.0038* | 0.0028* | 0.0025* | - |

**Table S4** - Whole genome sequencing reads produced and retained after filter-trimming.

| Year | Sample | Read Pairs Produced | Pairs surviving Filter-Trimming |
|---|---|---|---|
| 2002 | AMD_2002_1_S1 | 17857972 | 15183065 |
| 2002 | AMD_2002_2_S2 | 18328697 | 15496427 |
| 2002 | AMD_2002_3_S3 | 17308628 | 14708405 |
| 2002 | AMD_2002_4_S4 | 17233054 | 14498291 |
| 2002 | AMD_2002_5_S5 | 18353215 | 15540410 |
| 2002 | AMD_2002_6_S6 | 19357678 | 16478819 |
| 2002 | AMD_2002_7_S7 | 17653925 | 14891780 |
| 2002 | AMD_2002_8_S8 | 20942008 | 17706839 |
| 2002 | AMD_2002_9_S9 | 17545784 | 14895669 |
| 2002 | AMD_2002_10_S10 | 16907357 | 13234484 |
| 2002 | AMD_2002_11_S11 | 21096016 | 17897536 |
| 2002 | AMD_2002_12_S12 | 19119251 | 16056352 |
| 2002 | AMD_2002_13_S13 | 16686090 | 14112995 |
| 2012 | 2012_1_S17 | 14220211 | 11333084 |
| 2012 | 2012_2_S18 | 14838561 | 11301819 |

| | | | |
|---|---|---|---|
| 2012 | 2012_3_S19 | 16022353 | 12620146 |
| 2012 | 2012_4_S20 | 11837225 | 9140148 |
| 2012 | 2012_5_S21 | 12770603 | 10036611 |
| 2012 | 2012_6_S22 | 14175355 | 11205330 |
| 2012 | 2012_7_S23 | 13877872 | 10820989 |
| 2012 | AMD_2012_9_S14 | 16993947 | 14228141 |
| 2012 | AMD_2012_10_S15 | 19183909 | 16133697 |
| 2012 | AMD_2012_11_S16 | 16334186 | 13534648 |
| 2012 | AMD_2012_12_S17 | 19214481 | 15890092 |
| 2017 | 2017_1_S25 | 13823437 | 11111671 |
| 2017 | 2017_3_S26 | 14570254 | 11586632 |
| 2017 | 2017_5_S27 | 14256089 | 11458471 |
| 2017 | 2017_6_S28 | 14216234 | 11216536 |
| 2017 | 2017_7_S29 | 14271707 | 11367216 |
| 2017 | 2017_8_S30 | 14902242 | 11992610 |
| 2017 | 2017_9_S31 | 14121221 | 11268287 |
| 2017 | AMD_2017_9_S19 | 18482153 | 15466675 |

| | | | |
|---|---|---|---|
| 2017 | AMD_2017_10_S20 | 19547412 | 16356430 |
| 2017 | AMD_2017_11_S21 | 19857182 | 16543114 |
| 2017 | AMD_2017_12_S22 | 17291804 | 14526909 |

**Table S5 -** Probabilities of genomic divergence equivalent to those seen in WGS experiments under neutral conditions with varying present day population sizes ($N_0$). Threshold values were generated from our WGS experiments, and include our empirically-derived $ZF_{ST}$ threshold of greater than 6 ($F_{ST} = 0.047$), and the two most extreme values identified for 40kb windows on scaffolds 569 ($F_{ST} = 0.089$) and 1612 ($F_{ST} = 0.146$).

| **$N_0$** | **Total Sims** | **Mean $F_{ST}$** | **prob $F_{ST}$ > 0.047** | **prob $F_{ST}$ > 0.089** | **prob $F_{ST}$ > 0.146** |
|---|---|---|---|---|---|
| 21,000 | 20,000 | 0.00061 | 0.0189 | 0 | 0 |
| 63,000 | 20,000 | 0.00010 | 0.0005 | 0 | 0 |
| 105,000 | 20,000 | 0.00015 | 0.0001 | 0 | 0 |
| 140,000 | 20,000 | 0.0001 | 0 | 0 | 0 |

**Table S6** - Genes from families known to be associated with detoxification or Cry resistance within 50 kb of regions with higher than expected temporal genetic divergence in *H. zea* from Bossier Parish, LA.

| Gene Family | Gene ID | Gene annotation |
|---|---|---|
| ABC transporter | HzOG200336 | *abcB6* |
| choline/carboxylesterase | HzOG200194 | *cce001e* |
| | HzOG200193 | *cce001m* |
| | HzOG200152 | *cce024b* |
| | HzOG200151 | *cce024a* |
| cytochrome p450 | HzOG200024 | *cyp333b3* |
| | HzOG200001 | *cyp15c1* |
| peptidase | HzOG204148 | *venom dipeptidyl peptidase 4-like* |
| | HzOG204151 | *carboxypeptidase Q-like* |
| | HzOG204153 | *carboxypeptidase Q-like* |
| tetraspanin | HzOG211510 | *tetraspanin 68C* |

**Table S7 -** Allele frequencies at the sites of non-synonymous SNP substitutions in the LA (n = 22) and MD populations (n = 16 and 16, respectively) collected in 2017. We compared allele frequencies compared using a Freeman-Halton extension of a Fisher's exact test with a Bonferroni correction ($\alpha$ = 0.006) as a test of statistical significance.

| **Scaf** | **Gene** | **SNP Pos (bp)** | **Ref Codon** | **LA 2017 freq** | **MD 2017 Bt freq** | **MD 2017 Non-Bt freq** | **p-val** |
|---|---|---|---|---|---|---|---|
| KZ118395.1 | *cpq1* | 9,409 | GCT | 0.14 | 0.19 | 0 | 0.23 |
| KZ118395.1 | *cpq2* | 152,183 | ATA | 1 | 0.81 | 0.87 | 0.10 |
| KZ118395.1 | *cpq2* | 152,595 | GTA | 1 | 0.81 | 0.81 | 0.05 |
| KZ117131.1 | *cyp333b3* | 35,685 | AGC | 0.18 | 0.19 | 0 | 0.18 |
| KZ117131.1 | *cyp333b3* | 36,339 | TTT | 0.09 | 0.07 | 0.19 | 0.64 |
| KZ117131.1 | *cyp333b3* | 36,579 | GGA | 0.09 | 0.25 | 0 | 0.08 |
| KZ117131.1 | *cyp333b3* | 36,959 | GCC | 1 | 0.94 | 1 | 0.59 |
| KZ117131.1 | *cyp333b3* | 37,490 | AGC | 0.09 | 0.25 | 0 | 0.08 |

**Table S8** - PCR primers targeting non-synonymous mutations in *cpq1* and *cyp333b3*. Target sequences can be amplified using the following thermal cycler conditions: 2 min denaturation at 95C followed by 30 cycles of 95 C for 30 sec, 58 C for 30 sec, and 72 C for 40 sec.

| **Gene** | **Primer Pair** |
|---|---|
| *cpq1* | F: 5' - TTGAAGTCTTCTCATCAAATCTGC<br>R: 5' - GATGCACATTACGTGACTTATGG |
| *cyp333b3* | F: 5' - CCGTATCTACACCAGCGAATAA<br>R: 5' - CTGTTCAAGCAAGCCATGAAA |

**Table S9 –** Likelihood ratio test of Cry resistance linkage to *cpq1* and *cyp333b3*. The number of F2 individuals analyzed for each diet treatment is represented by n.

| **Gene** | **Diet** | **df** | **χ2** | **p-value** | **n** |
|---|---|---|---|---|---|
| *cpq1* | untreated | 2 | 23.6 | < 0.0001 | 50 |
| *cpq1* | Cry1Ab | 2 | 0.1 | 0.94 | 58 |
| *cpq1* | Cry1A.105+Cry2Ab2 | 2 | 5.02 | 0.08 | 65 |

**Supplemental Figures**

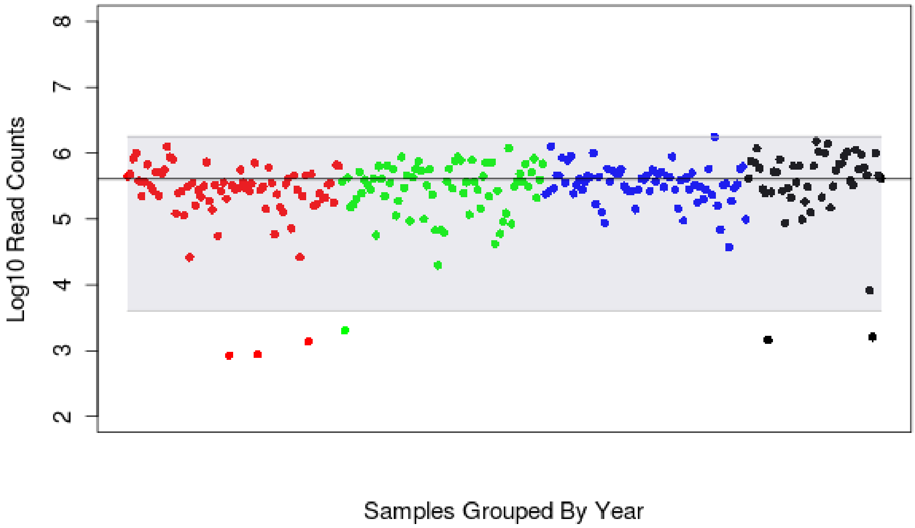


**Figure S1 -** $Log_{10}$ read count distribution for ddRAD-sequenced *H. zea* samples across years. Samples collected in 2002 (red), 2007 (green), 2012 (blue), and 2016 (black) are grouped by color along the x-axis. Those points that fell outside of the grey shaded region represent individuals whose read counts were lower than 4,000 (less than 1% of the mean read count for all libraries). Because of their low read counts, these 6 individuals were excluded from our population-level analyses.

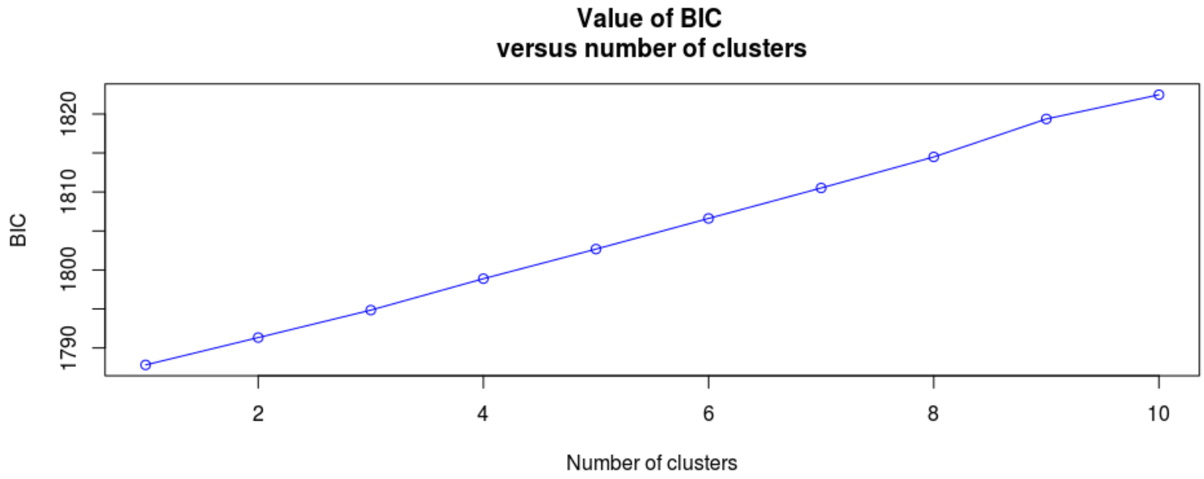


**Figure S2 -** K-means analysis, where model fit (BIC) is regressed on the number of putative populations (K) for *H. zea* collected from Bossier Parish, LA, in the years 2002, 2007, 2012, and 2016. This indicates that a single population was the most appropriate fit for our data.

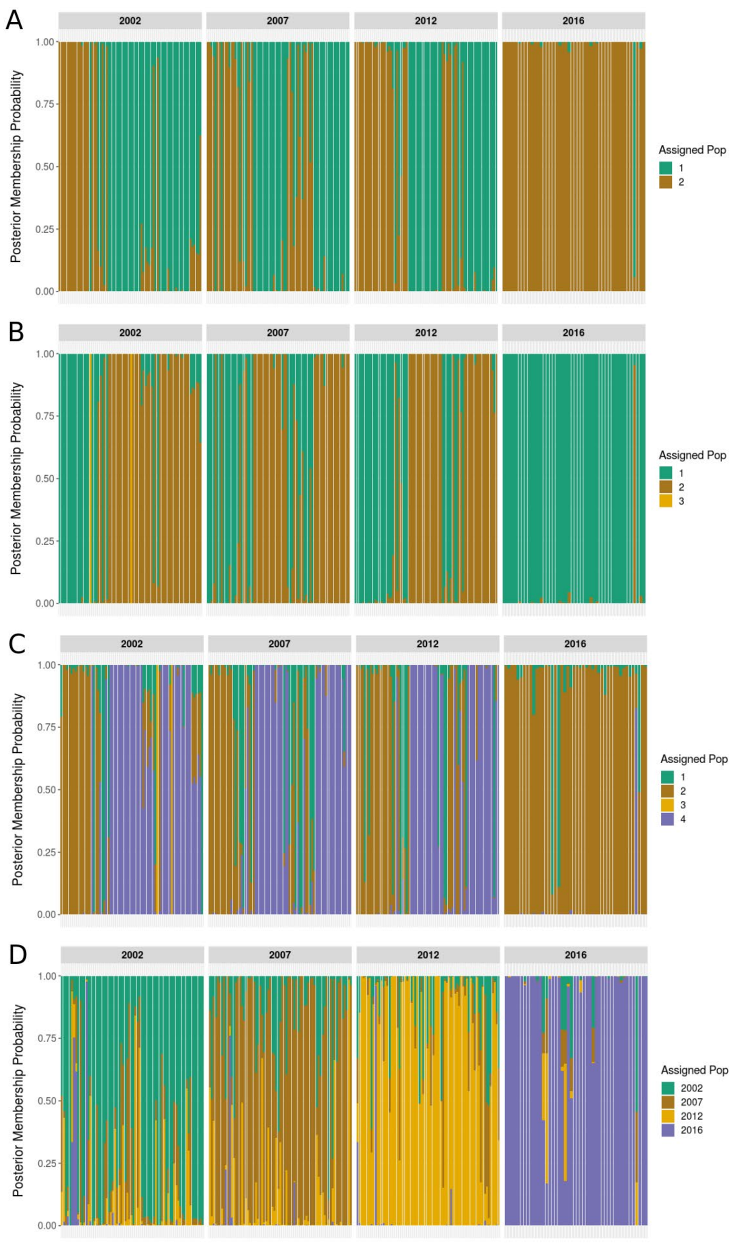


**Figure S3** - Posterior probabilities of population membership assigned to ddRAD-sequenced *H. zea* according to a discriminant analysis of principal components (DAPC). Each column represents a single individual, and individuals are grouped along the x-axis according to collection year. The analysis was conducted using the first 8 PCs and k = 2-4 inferred populations (panels A - C, respectively). Panel D shows posterior membership probabilities, where the original collection years were used as priors, along with the first 115 PCs.

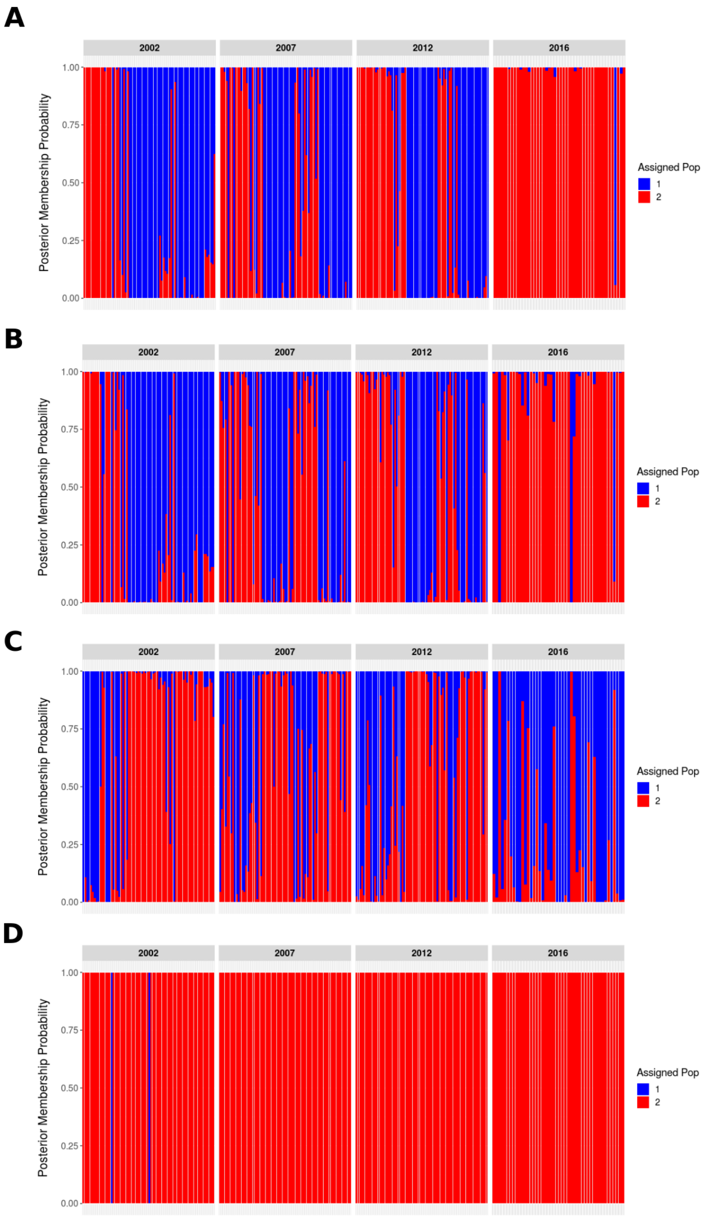


**Figure S4** - Posterior membership probabilities according to DAPC for ddRAD-sequenced *H. zea* assuming k = 2 clusters or groups. For this series of DAPCs, we removed the top 1-5% of variance contributing SNPs and then estimated posterior membership probability for each individual. Each individual is represented by a single column, and individuals from each collection year are grouped together along the x-axis. Numbers of SNPs used for each DAPC were (A) 14,398, (B) 14,254, (C) 14,038, and (D) 13,678.

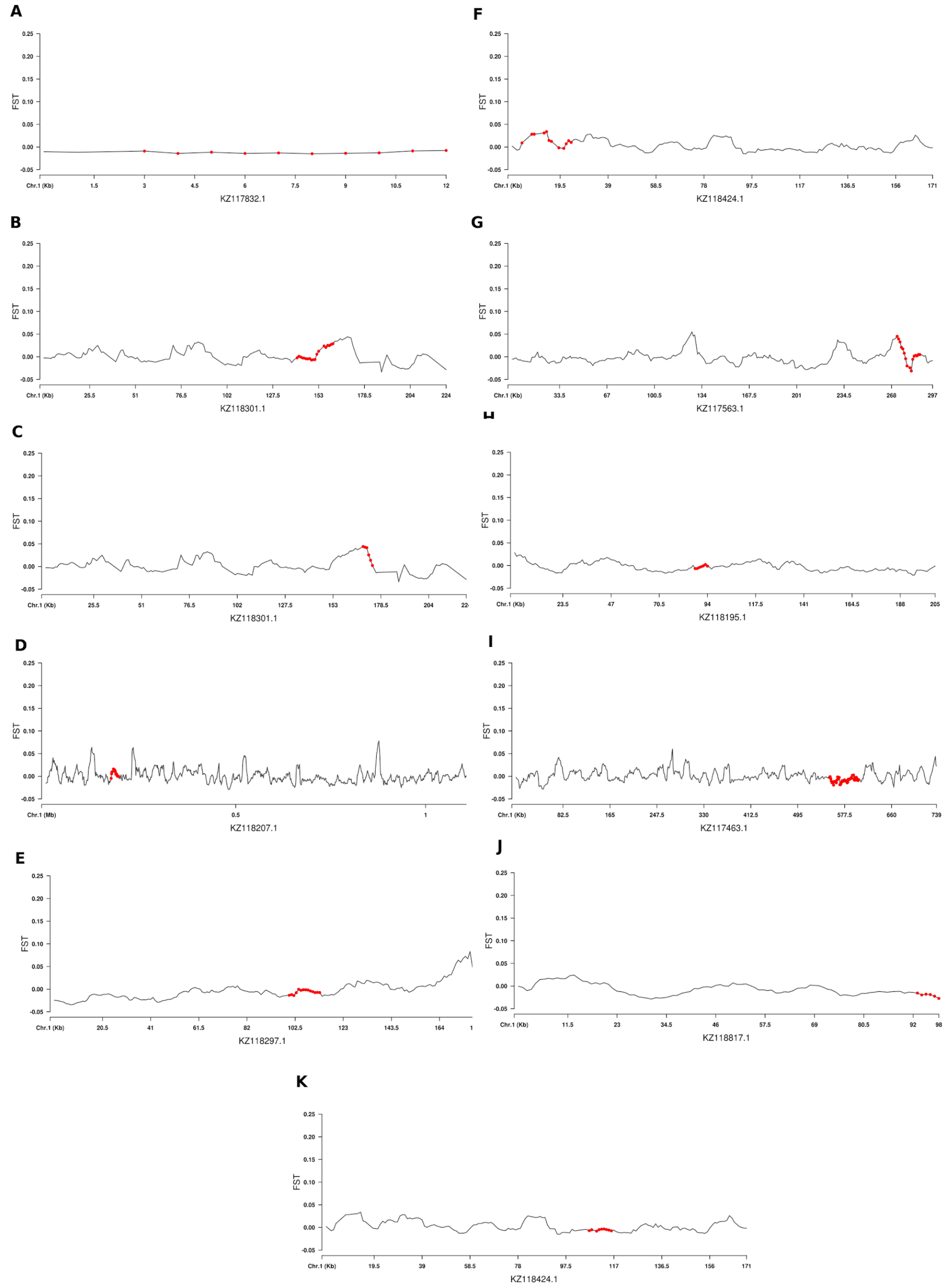


**Figure S5** - Average $F_{ST}$ values for 10kb genomic windows with a 1 kb step size along *H. zea* scaffolds containing previously described Bt resistance candidate genes: *alp* (A), *apn1* (B), *apn4* (C), *abcA2* (D), *abcC2* (E), *abcG1* (F), *calp* (G), *cad2* (H), *cad-86C* (I), *map4K4* (J), *tspan1* (K). Windows containing the genes are indicated in red. Scale of the x-axis varies by plot depending upon the length of the scaffold.

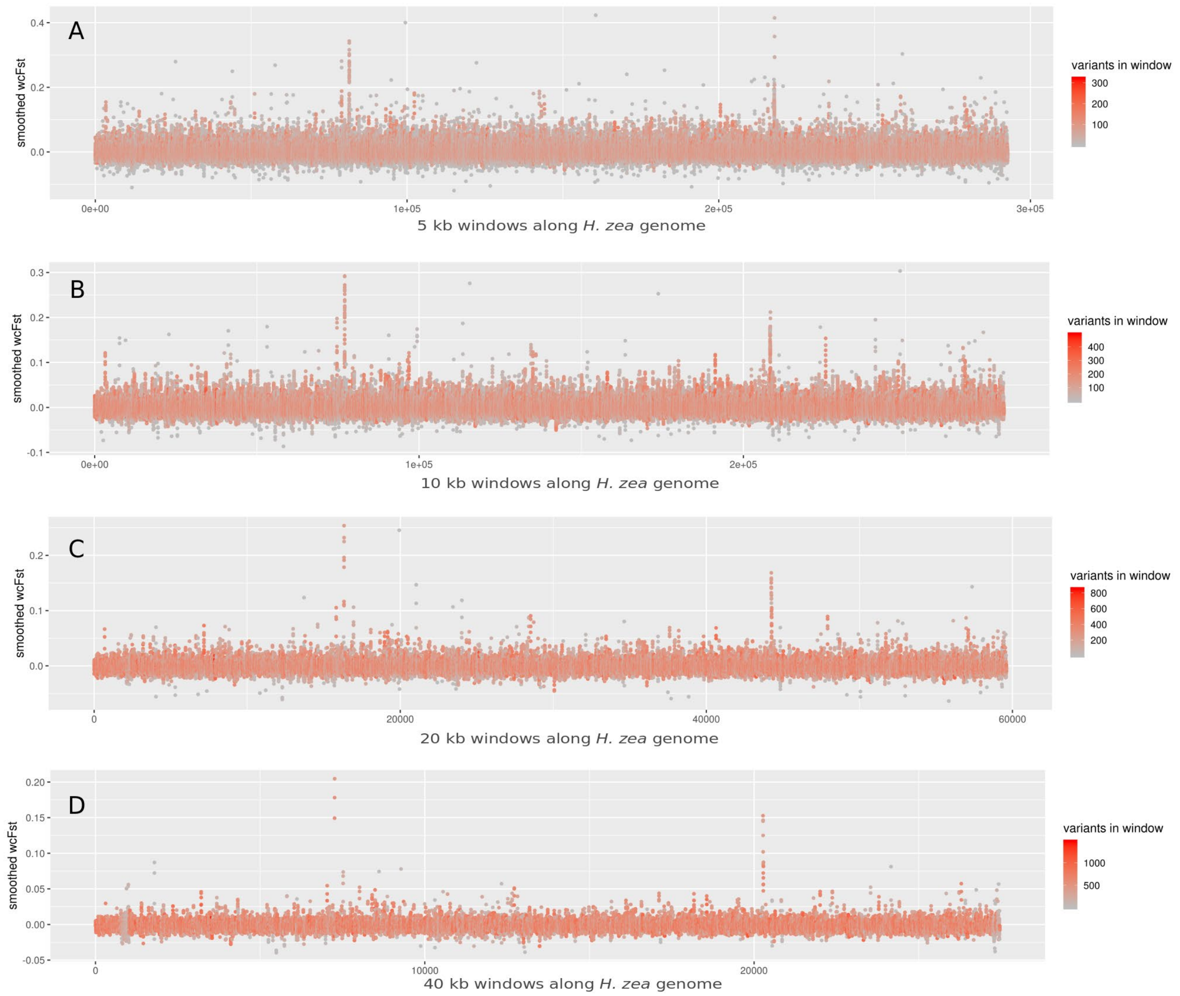


**Figure S6 -** Weir and Cockerham's averaged $F_{ST}$ as calculated for the 2002 and 2017 collections of *H. zea* samples that underwent whole genome sequencing. Sliding window/step sizes were A) 5/1 kb, B) 10/1 kb, C) 20/5 kb, and D) 40/10 kb. Point colors indicate the number of SNP variants used to calculate the sliding window averaged $F_{ST}$ values.

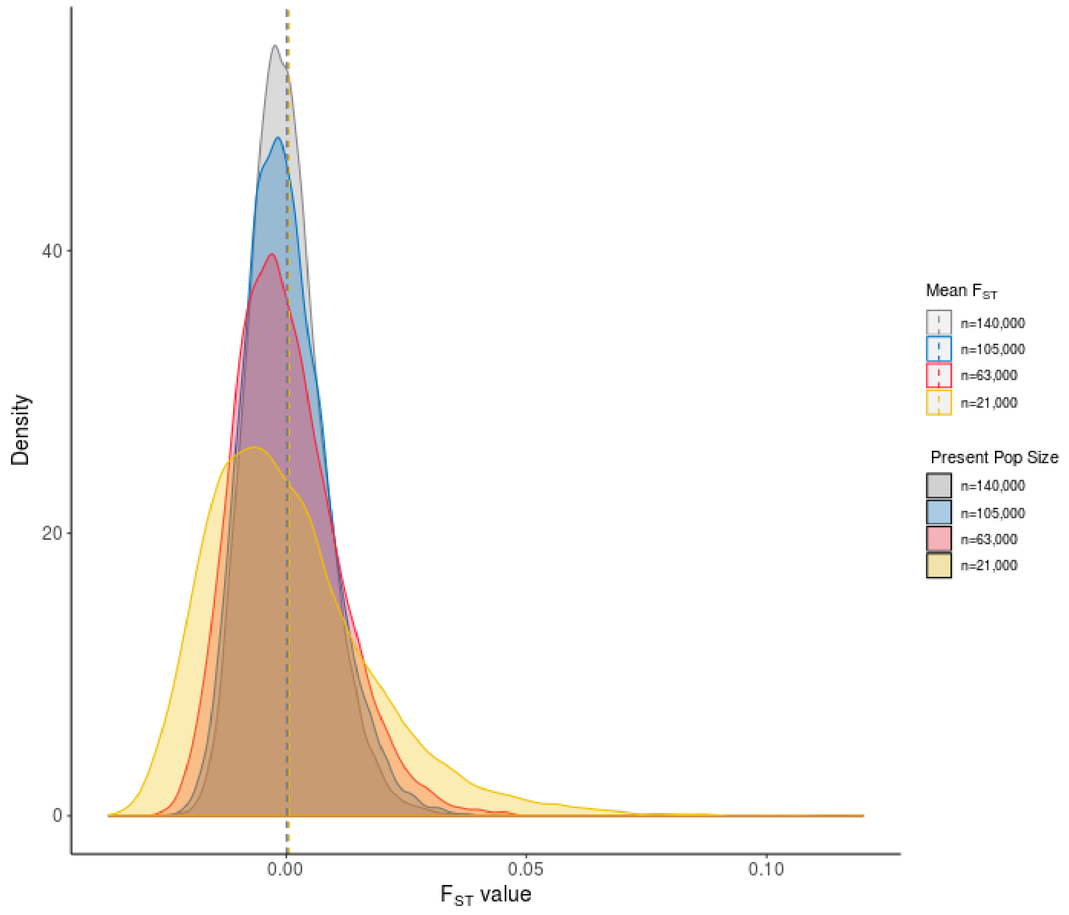


**Figure S7 -** Distributions of $F_{ST}$ values for ms simulated genotypes from an unselected "present day" population compared to the ancestral population 75 generations in the past. Present day population sizes varied from 21,000 to 140,000 (indicated by color), and we assumed a constant mutation rate of 2.9 x $10^{-9}$, and a recombination rate of 4cM/Mb.

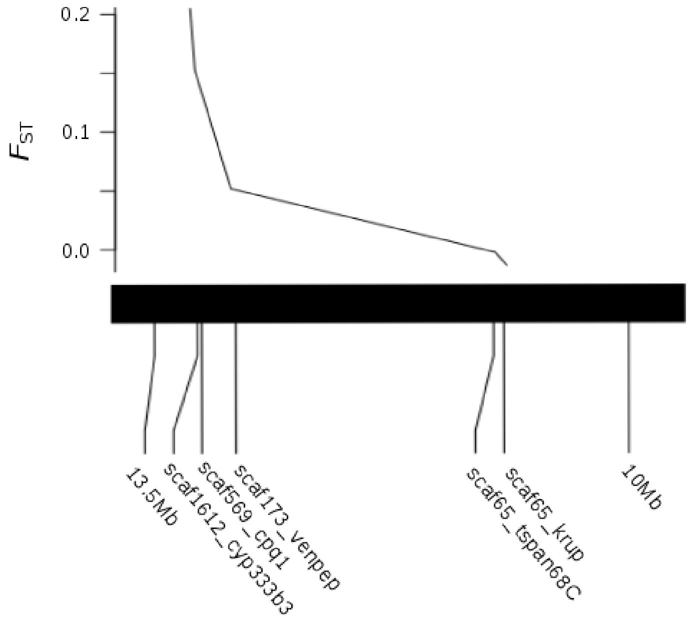


**Figure S8** - A tblastn alignment of genes on the most divergent *H. zea* scaffolds to the *B. mori* genome assembly revealed that 4 were physically linked and syntenic to *B. mori* chromosome 13. Forty kilobase window averaged $F_{ST}$ values from the 2002-2017 comparison near each gene is plotted above the linkage map. Scaffold 65 values were low for the 2002-2017 comparison but higher than expected in the 2002-2012 comparison ($F_{ST} \sim 0.05$). This indicated that this region experienced selection during the earlier time period, but divergence did not persist over time, likely due to the breakdown of linkage disequilibrium.

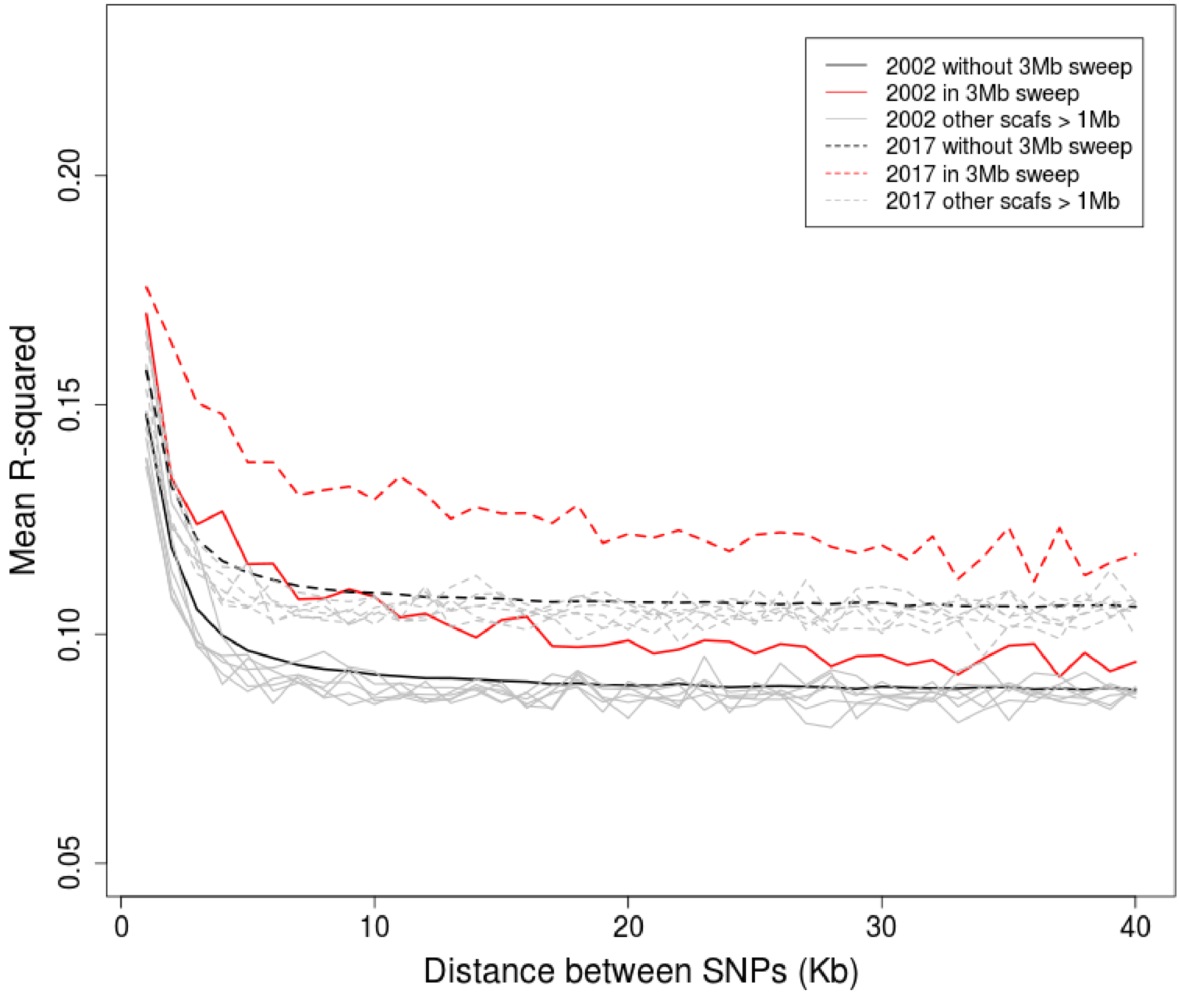


**Figure S9** - Decay of linkage disequilibrium within 40 Kb windows in the *H. zea* genome, as measured by the square of the correlation coefficient between allele frequencies.

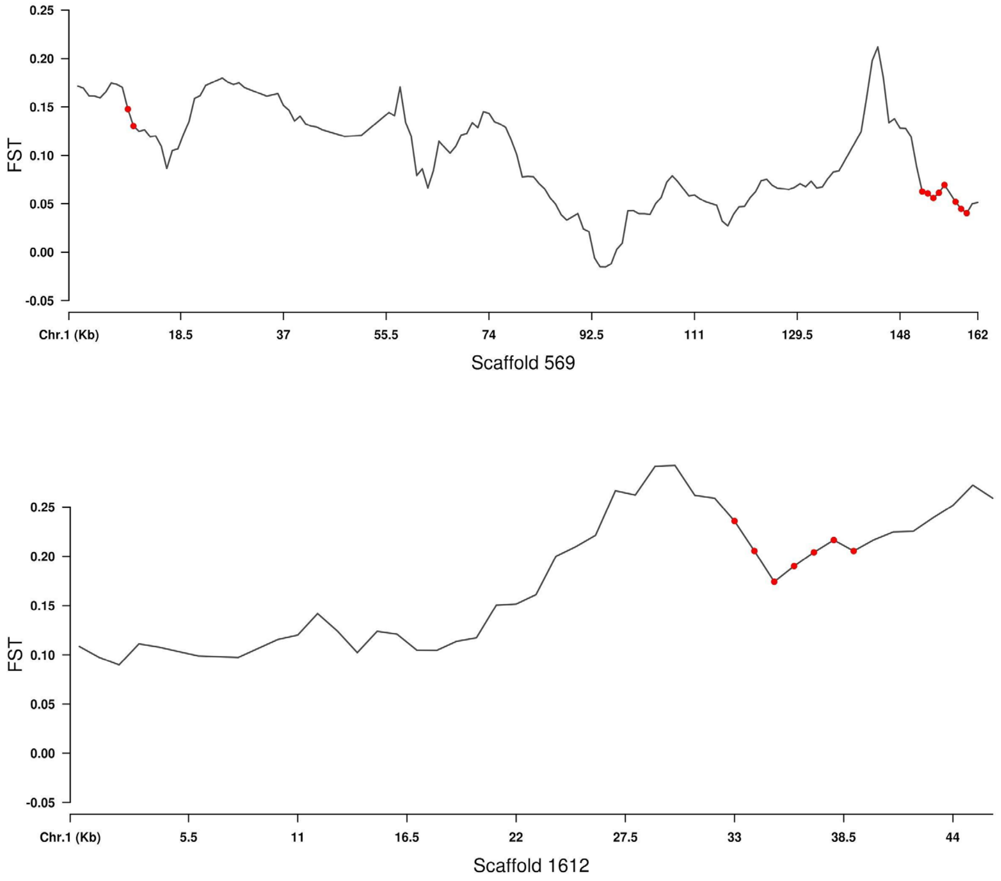


**Figure S10** - Ten kb sliding-window averaged $F_{ST}$ (1kb step size) showing genomic divergence along scaffolds 569 and 1612 between *H. zea* adult males collected in 2002 and 2017. The *cpq1* and *cpq2* are seen in red in the top panel, while the *cyp333b3* is in red in the bottom panel.

### Supplemental Methods

*Insect collections for ddRAD-seq and WGS analyses*

*H. zea* adults were collected by pheromone-baited trap bi-weekly from May through September in Bossier Parish, Louisiana (Table S1). Collections were made in the years 2002, 2007, 2012, 2016, and 2017, and specimens were stored in 100% ethanol at -20° C until use. Late instar (5-6$^{th}$) *H. zea* were also collected in August and September of 2017 at the University of Maryland CMREC farms in Prince George's county, MD, from the ears of Bt-expressing and non-Bt sweet corn isolines according to Dively *et al.* (2016) and reared to pupation under laboratory conditions of 16:8 LD at 25° C and 50% RH. Newly emerged adults were sacrificed by freezing and held at -80° C until use. Genomic DNA was isolated from half of an adult thorax with a Qiagen DNeasy kit (Qiagen, Inc., Valencia, CA, USA) using a modified mouse tail protocol according to Fritz et al. (2020).

*DdRAD-seq library preparation details*

Specimens from the years 2002, 2007, 2012, and 2016 were prepared into Double-digest RAD-seq (ddRAD-seq) libraries for population genomic analysis. Libraries were prepared according to Fritz *et al*. (2016; 2018). Briefly, genomic DNA was fragmented using EcoRI and MSPI, and DNA fragments from each individual were barcoded with unique 6-mer or 8-mer nucleotide sequences. Barcoded DNA from no more than 11 adults were pooled and size-selected for fragments ranging from 450-650 base-pairs (bp). A polymerase chain reaction (PCR) added a second identifier, a standard Illumina TruSeq index, such that reads from each specimen could be identified by their unique combination of barcode and index. Indexed pools were combined in equimolar amounts (4 nM) for sequencing submission to the North Carolina

State Genomic Sciences Laboratory. In total, we sequenced 265 individuals spread across 8 runs of an Illumina MiSeq. Individuals with low read counts (< 4000 reads; Figure S1) were excluded from our downstream population genomic analysis such that our final sample size was 259 individuals.

*ddRAD-seq Bioinformatic Analysis*

Adapter sequences were trimmed from Illumina reads using cutadapt (v. 1.18; Martin, 2011), and read pairs were merged to allow for a maximum overlap of 200 bp (FLASh v. 1.2.11; Magoc and Salzberg, 2011). Merged reads were demultiplexed and filtered using the process_radtags script from Stacks (v. 2.2; Catchen et al., 2011) to remove reads without an intact EcoRI cut site, or a quality score < 30. We did not allow process_radtags to rescue barcodes. Filter-trimmed reads were aligned to the *H. zea* genome assembly (v. 1.0, NCBI Bioproject PRJNA378438; Pearce et al., 2017) with Bowtie2 using the highest sensitivity settings in end-to-end mode (v. 2.2.6-2; Langmead and Salzberg, 2012).

Following genome alignment, we genotyped individuals using BCFtools (Danecek et al., 2016), where SNPs were called with the mpileup and call functions using standard options. We only used reads with mapping quality of 5 or higher for genotyping. Downstream of genotyping, our dataset was filtered by vcftools (Danecek *et al.,* 2011) to include only SNPs that i) had a depth of coverage > 3, ii) were present in at least 75% of the 259 individuals in our filtered dataset, iii) had a minor allele frequency of 0.05, iv) had a maximum of 2 alleles, and v) were thinned to 1 per 200 bp to reduce the degree of linkage disequilibrium between SNPs on the same ddRAD-seq marker.

*Overall population genomic divergence across years*

We calculated average nucleotide diversity ($\pi$) and heterozygosity (the inbreeding coefficient F) values and their corresponding 95% bootstrapped confidence intervals (N = 5000) for each population using vcftools. Weir and Cockerham's Pairwise $F_{ST}$ values (1984) were calculated to quantify overall population genomic divergence between years with the StAMPP package (v. 1.5.1, Pembleton *et al.,* 2013) in R (v. 3.4.4, R Development Core Team, 2008).

A k-means analysis was implemented with the R package adegenet (v. 2.1.1, Jombart and Ahmed, 2011), and results were compared those of a Pairwise $F_{ST}$ analysis. We retained 225 PCs and compared models containing 1:10 clusters using Bayesian Information Criterion (BIC). For the four best-fitting numbers of clusters (k = 2-5), we conducted a Discriminant Analysis of Principal Components (DAPC) with adegenet to determine whether there was significant clustering of individuals within and between years according to their ddRAD-seq genotypes. The xvalDapc function was used to cross-validate the numbers of principal components (PCs) required for accurate placement of individuals into clusters. We examined posterior membership probabilities for individuals both according to: 1) inferred clusters (k = 2-4) using the first 8 PCs as indicated by our cross-validation, as well as 2) clustering by collection year using the first 115 PCs.

We identified SNPs with significant allele frequency divergence over time with an outlier analysis using the R package OutFLANK (v. 0.2, Whitlock and Lotterhos 2015). Weir and Cockerham's $F_{ST}$ was calculated for each SNP to quantify genetic divergence across years. For analysis, our left- and right-trimming fractions were set to 0.05, the minimum heterozygosity value was 0.1, the false discovery rate threshold was 0.1, and $\alpha$ was specified as 0.05.

A number of genes have already been implicated in Bt resistance in other Lepidopteran pest species (Table 1). A blast search (https://blast.ncbi.nlm.nih.gov) identified the locations of these putative Bt resistance genes in the *H. zea* genome. Using our ddRAD-seq data, we first determined whether ddRAD-seq genomic outliers were found near or within any of the known 11 Bt resistance candidate genes. We also report the distance between the nearest ddRAD-seq marker and each of these genes (Table 1).

*WGS library preparation*

To confirm our ddRAD-seq results, and extend our analysis to regions not covered by ddRAD-seq markers, we used whole genome resequencing to scan the genomes of additional *H. zea* collected from this same location in LA. Samples of gDNA from additional adults collected in 2002 (n= 13), 2012 (n = 11), and 2017 (n = 11) were submitted to the North Carolina State University Genomic Sciences Laboratory for Illumina TruSeq LT library preparation (Illumina, Inc. San Diego, CA). DNA samples were barcoded for identification of each individual and subsequently pooled for sequencing. The prepared libraries were sequenced on two Illumina NextSeq500 runs at the North Carolina State University Genomic Sciences Laboratory using 150 bp paired-end reads.

*WGS bioinformatic analysis*

Reads were trimmed to remove Illumina TruSeq adapters and low quality sequence regions (5 bp averaged quality scores of < 20) and filtered to retain reads in proper pairs that were at least 50bp long (Trimmomatic v. 0.38; Bolger et al., 2014). Filter-trimmed reads were aligned to the *H. zea* reference genome with Bowtie2 as described for our ddRAD-seq analysis.

Reads with an alignment quality of < 5 were filtered out using SAMtools (v. 0.1.19; Li et al., 2009), and duplicate reads (PCR and optical) were removed using GATK (v. 4.0.12.0; McKenna et al. 2010). Filtered alignment files were converted to a Variant Call Format (vcf) file using BCFtools, and indels were removed with vcftools. We further filtered our SNPs to remove those i) with a coverage depth < 3, ii) absent from 8 or more individuals sequenced, iii) with minor allele frequency of < 0.05, and iv) with > 2 alleles per locus.

*Bioinformatic identification of genomic regions under selection*

We used vcftools to calculate sliding window averaged $F_{ST}$ values across the entire genome with a 40kb window and a 10kb step size for samples from each pair of years. Additional sliding window and step sizes were also examined for the comparison of samples collected in 2002 and 2017, and are reported in Figure S2. To prevent spurious signals from few SNPs, we excluded windows with fewer than 10. According to Rubin et al. (2010), we standardized estimates of $F_{ST}$ with a z-transformation, and windows with an average $F_{ST}$ value greater than 6 standard deviations from the genome-wide average $F_{ST}$ value ($ZF_{ST} > 6$) were considered to have undergone statistically significant genetic divergence. This value was calculated both for 40kb windows with a 10kb step size, and 10 kb windows with a 1kb step size. Genotype simulations under neutral expectations using realistic population demographic conditions generated a distribution of $F_{ST}$ values, which were compared against empirical $F_{ST}$ values from our genome scan results as a secondary approach to examining statistical significance (Hudson 2002; File S4).

We then examined the genes within 50kb of each genomic window showing signs of statistically significant genetic divergence between pairs of years (2002-2017, 2002-2012, 2012-

2017) using custom R scripts. We queried the gene list to identify those from families generally known to be associated with detoxification of plant-produced or human-applied toxicants using the following generic search terms: “peptidase”, “cce”, “protease”, “proteinase”, “cyp”, “cadherin”, “abc”, “phosphatase”, “transferase”, “voltage-gated”, “tetraspanin”, and “gst”, which matched the terms/abbreviations used for these gene families in the *H. zea* functional annotation (gff) file accessed through the CSIRO data access portal (https://doi.org/10.4225/08/598d49cb2cd23). Finally, we compared the results of our whole genome scan to those of our ddRAD-enabled outlier analysis to determine whether we could have arrived at the same conclusions using ddRAD-seq alone.

The genome assembly of *H. zea* is fragmented into 2,975 genomic scaffolds, which makes detecting the extent of a selective sweep challenging. We reasoned that under conditions of strong selection some of these scaffolds with elevated genomic divergence should be physically linked. Therefore, a tblastn alignment of genes from the 5 most divergent scaffolds to the *Bombyx mori* genome available through Kaikobase (v. 3.2.2; http://sgp.dna.affrc.go.jp/KAIKObase/) identified their relative positions using conserved macrosynteny among Lepidoptera (d'Alençon et al. 2009).

*ms simulations*

We simulated 40 kb genomic windows for individuals from each of two populations under conditions of no selection using Hudson’s ms (Hudson 2002). Weir and Cockerham’s $F_{ST}$ values were used to quantify genetic divergence between the two simulated populations, to quantify the probability that stochastic processes (e.g. genetic drift) could have produced $F_{ST}$ values as extreme as 0.047 ($ZF_{ST} > 6$ per 40kb), our cutoff for statistical significance in our genome scan. Genotypes from individuals sampled in the present day were compared with those from individuals sampled from the same population 75 generations (5 generations/year for 15 years; Reay-Jones 2019) in the past. We conducted 20,000 simulations, using a population demographic model that assumed a mutation rate of 2.9 x $10^{-9}$ (Anderson et al. 2018), and a recombination rate of 4cM/Mb (Martin et al. 2019). Present day population sizes varied as follows: n = 21,000, 63,000, 105,000, and 140,000. These assumed a trapping radius of 1 acre (Lopez and Witz 1988), a corn seeding rate of 28,000 per 5000 $m^2$ (Morris et al. 2000), an infestation rate of 50-100% (Dively et al. 2016), and a rate of larval survivorship of 30-100% (Pan et al. 2016). The product of the seeding rate per acre, infestation rate, and larval survivorship rate gives the estimated population size within a 1 acre trapping area, which we multiplied by 5 to account for our 5 traps (Table S1). As in our WGS genome scan, we verified that all simulations produced at least 10 segregating sites prior to calculating $F_{ST}$ with the hierfstat package (v. 0.04-22; Goudet 2005) in R. Distributions of $F_{ST}$ values generated for each simulated experiment with varying present day population sizes are presented in Table S5 and were compared with empirically derived $F_{ST}$ values from our WGS experiments.

*Genome similarity between* H. zea *collected from MD and LA in 2017*

A global $F_{ST}$ analysis was implemented using the R package hierfstat (v. 0.04-22; Goudet 2005), and compared *H. zea* collected at CMREC in Prince George's county, MD, in 2017 to those collected in Bossier Parish, LA, in 2017. Bootstrapped 95% confidence intervals around the global $F_{ST}$ value were generated using the R package bootstrap (v. 2019.6; Tibshirani and Efron 1993). Overlap of the 95% confidence intervals with zero indicated a lack of statistical significance.

*Gene-specific analyses*

Putative protein-coding changes were examined for three genes found within regions of highest genetic divergence across years, *cpq1*, *cpq2* and *cyp333b3*. We extracted the nucleotide sequence for the coding region of each gene using a custom bioinformatic pipeline, and ExPASy (https://web.expasy.org/translate/) was used to produce the putative protein sequences from individuals collected in 2002 and 2017. Protein sequences were aligned to one another using MUSCLE (Edgar 2004) to identify putative amino acid substitutions. Allele frequencies for SNPs which caused these substitutions were calculated for each collection year. A two-sided 3x2 Freeman-Halton extension of the Fisher's exact tests with a Bonferroni-corrected alpha value ($\alpha$ = 0.006) was used to examine whether allele frequencies differed significantly among years. Frequencies of these 8 non-synonymous SNPs were also compared in 2017 collections of *H. zea* from MD and LA using this same approach to quantify the extent of similarity over geographical distance.

*Strength of selection*

The strength of selection ($s$) imposed upon ancestral codons in the protein-coding regions of *cpq1*, *cpq2*, and *cyp333b3* were examined over time (Falconer and Mackay 1996). We calculated $s$ for two periods of selection: 2002 - 2012, as well as 2012 - 2017. We assumed 5 generations per year (Reay-Jones 2019) for each selection period, and calculations were performed assuming dominance of the allele that was increasing in frequency, $p$, no dominance of $p$, and recessiveness of $p$.

*Frequency of non-synonymous substitutions prior to 2002*

Genomic DNA was isolated from *H. zea* collected in 1998 from Hartstack traps in Bossier Parish, LA, 1998 using a Qiagen DNeasy isolation kit as indicated above. DNA from each individual was PCR amplified using primers flanking each non-synonymous mutation (Table S8), which allowed us to determine the frequency of these mutations prior to 2002. Amplicons were sequenced on an ABI3730xl (Applied Biosystems, Foster City, CA, USA), and chromatograms were visually inspected for quality filtering and sample genotyping.

*Cry-treated bioassays and tests for linkage*

In 2019, we collected Cry resistant late instar (5-6$^{th}$) *H. zea* larvae from the ears of a Cry1A.105+Cry2Ab2 sweet corn hybrid (Performance Series® Obsession II, Seminis Vegetable Seeds, Inc.) and reared them to pupation on Southland's *H. zea* diet in the laboratory. Newly emerged field-collected adults were mated in single pairs to one another and the larval growth phenotypes of their progeny were bioassayed according to Dively et al. (2016). Briefly, finely ground lyophilized sweet corn leaf tissue from Cry expressing hybrids (Cry1Ab: Attribute® BC 0805, Cry1A.105+Cry2Ab2: Performance Series® Obsession II) and their non-Bt isolines (Cry1Ab: Providence, Cry1A.105+Cry2Ab2: Obsession) were combined at a diagnostic dose that significantly reduced growth in Cry susceptible *H. zea* (Benzon Research Inc., Carlisle, PA) and added to meridic diet (Southland's *H. zea* diet). We made both a single toxin (Cry1Ab only) treatment and a two toxin (Cry1A.105+Cry2Ab2) diet treatment, where the diagnostic doses were 6.4mg Cry1Ab leaf tissue per mL diet (equivalent to Dively et al. 2016 160 mg treatment), and 2.6 mg Cry1A.105+Cry2Ab2 leaf tissue per mL diet. Offspring from field collected adults (apprx. late 1$^{st}$ instar) were assayed on both Cry treated diet and corresponding untreated controls that contained an equivalent amount of non-Bt corn leaf tissue and 7 day growth was measured. Benzon larvae were assayed alongside field larvae to demonstrate differences in susceptibility between lines. We analyzed differences in the final weights of Benzon larvae and the progeny of MD field-collections using a model reduction approach, where a pair of generalized linear models with and without the effect of population (Benzon vs. MD Field) were compared by likelihood ratio test. Growth on each of the two diet types was analyzed separately, and because each of these datasets did not meet assumptions of normality, our models assumed a Gamma distribution of residuals.

Larvae from field populations whose 7 day growth on Cry treated diet indicated a resistant phenotype were transferred to Southland's diet to complete development and adults were crossed to Cry susceptible Benzon *H. zea* in an F2 genetic cross design. Two families of F2s were generated: 1) The first family exposed to Cry1Ab-treated diet had a resistant parent with a 7 day larval weight of 145.6 mg, which was 7.3 times greater than the mean 7 day larval weights for the Benzon susceptible strain. 2) A second family exposed to the Cry1A.105+Cry2Ab2 treated diet had a resistant parent with a 7 day larval weight of 149.2 mg, which was 23.0 times greater than that of the Benzon mean 7 day larval weight. Growth for half of the F2s from each family was measured on a diagnostic dose of tissue (Figure 5D & E) alongside Benzon controls, to ensure potency of the treatment. For the family tested on Cry1Ab-treated diet, half of the F2 progeny were placed on diet untreated leaf tissue from the non-Bt expressing isoline in equal concentration to the treated diet so that the effect of alleles important for speed of larval growth could be separated from those related to Cry resistance (Figure 5C). After phenotyping, F2s were genotyped at non-synonymous mutations in the *cpq1* and *cyp333b3* genes. For genotyping, DNA from each individual was PCR amplified using primers in Table

S8. Amplicons were sequenced on an ABI3730 (Applied Biosystems, Foster City, CA, USA) and genotypes were called using chromatogram files with the SangeranalyseR package (v. 0.99.25; Chao et al. 2020). We used a Kendall's tau correlation test to examine correlations between genotypes at these two loci within individuals. Due to the strength of the correlation (tau = 0.98; $p < 0.0001$), we conducted the linkage analysis using only the *cpq1* genotypes.

Associations between *cpq1* genotype and larval growth phenotype were analyzed for each set of F2 progeny on each diet type (untreated, Cry1Ab-treated, and Cry1A.105 + Cry2Ab2-treated) by linear mixed models using the R package lme4 (Bates et al. 2015). Full models assumed a gaussian distribution of residuals and included a fixed effect of genotype, as well as a random effect of diet batch (25mL meridic diet + leaf tissue mixture). To determine whether genotype explained a statistically significant amount of variation in 7 day larval weight, full models were compared to a reduced model without the effect of genotype using a likelihood ratio test. For F2 progeny on Cry1Ab-treated diet, 7 day larval weights were natural log transformed to meet assumptions of normality prior to modeling. For F2 progeny on Cry1A.105 + Cry2Ab2-treated diet, no transformation was needed, but we did analyze 7 day larval weights using two different full and reduced models. One full model specified diet batch as a random effect as described above, but due to an lme4 convergence warning, a second full model included diet batch as a fixed effect. Although the p-value was higher for this second model, we took a more conservative approach and report results from this second model.
